# Orbitofrontal high-gamma reflects spike-dissociable value and decision mechanisms

**DOI:** 10.1101/2024.04.02.587758

**Authors:** Dixit Sharma, Shira M. Lupkin, Vincent B. McGinty

## Abstract

The orbitofrontal cortex (OFC) plays a crucial role in value-based decision-making. While previous research has focused on spiking activity in OFC neurons, the role of OFC local field potentials (LFPs) in decision-making remains unclear. LFPs are important because they can reflect synaptic and subthreshold activity not directly coupled to spiking, and because they are potential targets for less invasive forms of brain-machine interface (BMI). We recorded LFPs and spiking activity using multi-channel vertical probes while monkeys performed a two-option value-based decision-making task. We compared the value- and decision-coding properties of high-gamma range LFPs (HG, 50-150 Hz) to the coding properties of spiking multi-unit activity (MUA) recorded concurrently on the same electrodes. Results show that HG and MUA both represent the values of decision targets, and that their representations have similar temporal profiles in a trial. However, we also identified value-coding properties of HG that were dissociable from the concurrently-measured MUA. On average across channels, HG amplitude increased monotonically with value, whereas the average value encoding in MUA was net neutral. HG also encoded a signal consistent with a comparison between the values of the two targets, a signal which was much weaker in MUA. In individual channels, HG was better able to predict choice outcomes than MUA; however, when simultaneously recorded channels were combined in population-based decoder, MUA provided more accurate predictions than HG. Interestingly, HG value representations were accentuated in channels in or near shallow cortical layers, suggesting a dissociation between neuronal sources of HG and MUA. In summary, we find that HG signals are dissociable from MUA with respect to cognitive variables encoded in prefrontal cortex, evident in the monotonic encoding of value, stronger encoding of value comparisons, and more accurate predictions about behavior. High-frequency LFPs may therefore be a viable – or even preferable – target for BMIs to assist cognitive function, opening the possibility for less invasive access to mental contents that would otherwise be observable only with spike-based measures.

## INTRODUCTION

Neural representations of value are essential for decision-making, learning, and other cognitive abilities. For decision-making in particular, converging evidence points to an important role for value representations in the orbitofrontal cortex (OFC). This includes evidence from electrophysiology (Padoa-Schioppa & Assad, 2006; L. Tremblay & Schultz, 1999), computational modeling (Rustichini & Padoa-Schioppa, 2015), human and animal lesion studies (Murray et al., 2015; Vaidya et al., 2018), and microstimulation (Ballesta et al., 2020; Knudsen & Wallis, 2020).

Much of what is known about the computations performed by OFC comes from examining spiking activity in single neurons. Many OFC neurons represent the value of items offered or chosen during decision-making (Hunt et al., 2018; Padoa-Schioppa & Assad, 2006; Thorpe et al., 1983; L. Tremblay & Schultz, 1999), and value representations defined by multi-neuron patterns of OFC firing can predict the outcome of value-based choices (McGinty & Lupkin, 2023; Rich & Wallis, 2016).

In comparison, far less is known about value- and decision-related information encoded by OFC local field potentials (LFPs). LFPs are an important electrophysiological signal because they reflect not only somatic and axonal currents generated by action potentials, but also the otherwise unobservable axon terminal and dendritic currents resulting from local synaptic transmission (Buzsáki et al., 2012; Einevoll et al., 2013). Thus, LFPs may furnish information about decision-related neural activity that is not observable from spiking in single neurons. In addition, LFPs represent potentially important targets for brain-machine interfaces (BMIs), which for practical reasons cannot always resolve spiking and must therefore rely on electrophysiological signals with lower spatial resolution (Stavisky et al., 2015).

LFPs in the high-gamma band (HG, typically 50 to 200 Hz) are a particularly attractive target for accessing neural processing without measuring spikes. This is because HG magnitude is often strongly correlated with the spiking of nearby neurons (Ray et al., 2008; Ray & Maunsell, 2011), and also reflects nearby synaptic currents and other subthreshold phenomena (Lindén et al., 2011; Reimann et al., 2013). This suggests the possibility that HG signals alone could be as – or more – informative than spiking signals for probing neural mechanisms, despite their lower spatial resolution. For example, Lundqvist et al. (2016, 2018) identified gamma-range LFP events, not reflecting local spiking, that encoded information in a working memory task.

Consistent with this view, Rich & Wallis (2017) showed that OFC spiking activity and concurrently measured HG signals both encoded the value of an anticipated reward in a non-decision-making operant task. However, they also identified differences between spiking and HG signals, in the form of distinct patterns of value encoding, defined by recording location and encoding latency. This dissociation between OFC spiking and HG value coding in a non-decision task suggests that a similar dissociation may also occur in a more complex decision-making context. Therefore, the goal of this study was to identify the unique properties of OFC HG and spiking during decision-making – particularly in terms of the representation and comparison of multiple decision options (Hunt et al., 2018; Strait et al., 2014).

To address this question, we measured the modulation of OFC HG by the values of items offered in each trial of a two-alternative forced-choice decision task. We then compared three properties of these HG value representations to those of concurrently measured spiking signals. First, we examined the magnitude, time course and sign of the modulation of HG as a function of the value of the target items. Such representations are thought to entail the input stage of a goods-based decision process (Padoa-Schioppa, 2011). Second, we asked how the representations of the values of the two targets were distributed across the population, especially the extent to which single HG channels simultaneously encoded both target values (Strait et al., 2014). Third, we quantified the choice-predictive accuracy of HG value signals, both in single channels and at the population level, and compared this to spike-based choice predictions. For all of these properties, we asked whether the HG signals merely reflected the information available from concurrently observed spiking or whether they departed from the properties of the spike signal.

The results show that compared to spiking, HG signals recorded on the same channels are more likely to be positively modulated by value, are more likely to reflect signatures of value comparison (Strait et al., 2014), and can furnish more accurate predictions about trial-by-trial choice outcomes. These findings corroborate the idea that HG in OFC does not merely reflect local spiking events (Ray et al., 2008), and suggest that value- and choice-relevant computations may be taking place at the level of synaptic interactions. In addition, these findings suggest that HG is a viable target for cognitive BMIs designed to monitor internal representations of value and choice intent and, for some applications, may furnish more informative signals than would be obtained from spiking alone.

## METHODS

### Data source

The neural data in this study (38 sessions, 26,318 total trials) includes a portion of the data reported previously in McGinty & Lupkin, (2023) (28 sessions, 19,936 total trials), as well as data not used in the previous publication (10 sessions, 6,382 total trials). The neural data analyses in McGinty & Lupkin, (2023) use sorted single units, whereas the present study uses multiunit activity (MUA) and HG signals obtained from LFPs. Some MUA-based results will therefore be similar (but not identical) to results reported by McGinty & Lupkin (2023); HG results are entirely new.

### Subjects and Apparatus

All experimental procedures were performed according to the NIH Guide for the Care and Use of Laboratory Animals and were approved by the Animal Care and Use Committees of Stanford University and Rutgers University – Newark. The study subjects were two adult male rhesus macaques, referred to as monkeys K and C, weighing ∼14 kgs each at the time of the study. Data from monkey C were acquired at Rutgers University – Newark, and data from Monkey K were acquired at Stanford University. The monkeys were implanted with MR-compatible head holders and recording chambers (Crist Instruments, Hagerstown, MD), and craniotomies were performed to access the orbitofrontal cortex. Monkey K had bilateral chambers allowing recording from both hemispheres, and Monkey C had a single chamber on the left hemisphere. All the surgical procedures were performed under isoflurane anesthesia using fully aseptic techniques and instruments. Analgesics and antibiotics were given pre-, intra-, and post-operatively as required. Both the monkeys had a minimum of 4 weeks of recovery time post-surgery.

Neurophysiology data were collected while monkeys were head-restrained and seated in a custom-built chair, with their eyes 57 cm away from a CRT monitor displaying the task stimuli (120Hz refresh rate, 1024×768 resolution). Three response levers (ENV-612M, Med Associates, Inc., St. Albans, VT) were placed in front of the subjects within their reach. One lever was placed 21 cm below the display center, and the other two levers were located ∼8.5 cm to the left and right of the center lever. Stimulus presentation, reward delivery, and monitoring of lever presses and eye position, were controlled through a set of custom scripts written by R. Kiani for the MATLAB computing environment (Mathworks, Inc., Natick, MA) and Psychtoolbox-3 (Kleiner et al., 2007). Eye movements were recorded non-invasively at a sampling rate of 250 Hz (Eyelink, SR Research, Mississauga, Ontario, Canada). Juice rewards were delivered via a gravity-fed reservoir and solenoid valve. Neural activity, eye movement, and task event data were acquired and stored using a Plexon Omniplex system (Plexon, Inc. Dallas, TX). Analyses were performed using custom code and standard toolboxes in MATLAB 2019b.

### Behavioral task

A condensed behavioral task sequence is illustrated in Fig. 1a. The complete task sequence is described as follows. Monkeys initiated a trial by fixating on a central point on the display and manually depressing the center lever. After holding fixation and simultaneously pressing the center lever for a variable duration of 1-1.5s, the fixation point disappeared, and two target arrays appeared, centered 7.5 degrees of visual angle to the left and right of the display center. Once the arrays appeared, the monkeys were allowed to shift their gaze to view the targets. Targets were initially masked and became visible only when the monkeys started moving their gaze toward one of the targets (see below). They were allowed to view the targets in any order and for any amount of time until they initiated their choice. The monkeys initiated their choice by lifting their hand from the center lever, and then indicated their choice by pressing the left or right lever within a 400ms (Monkey K) or 500ms deadline (Monkey C). This deadline discouraged monkeys from deliberating (e.g., changing their mind) after lifting their hand from the center lever. After the left/right lever press, a reward of 1-5 drops of juice was delivered according to the value of the chosen target (Fig. 1b).

**Fig. 1.**
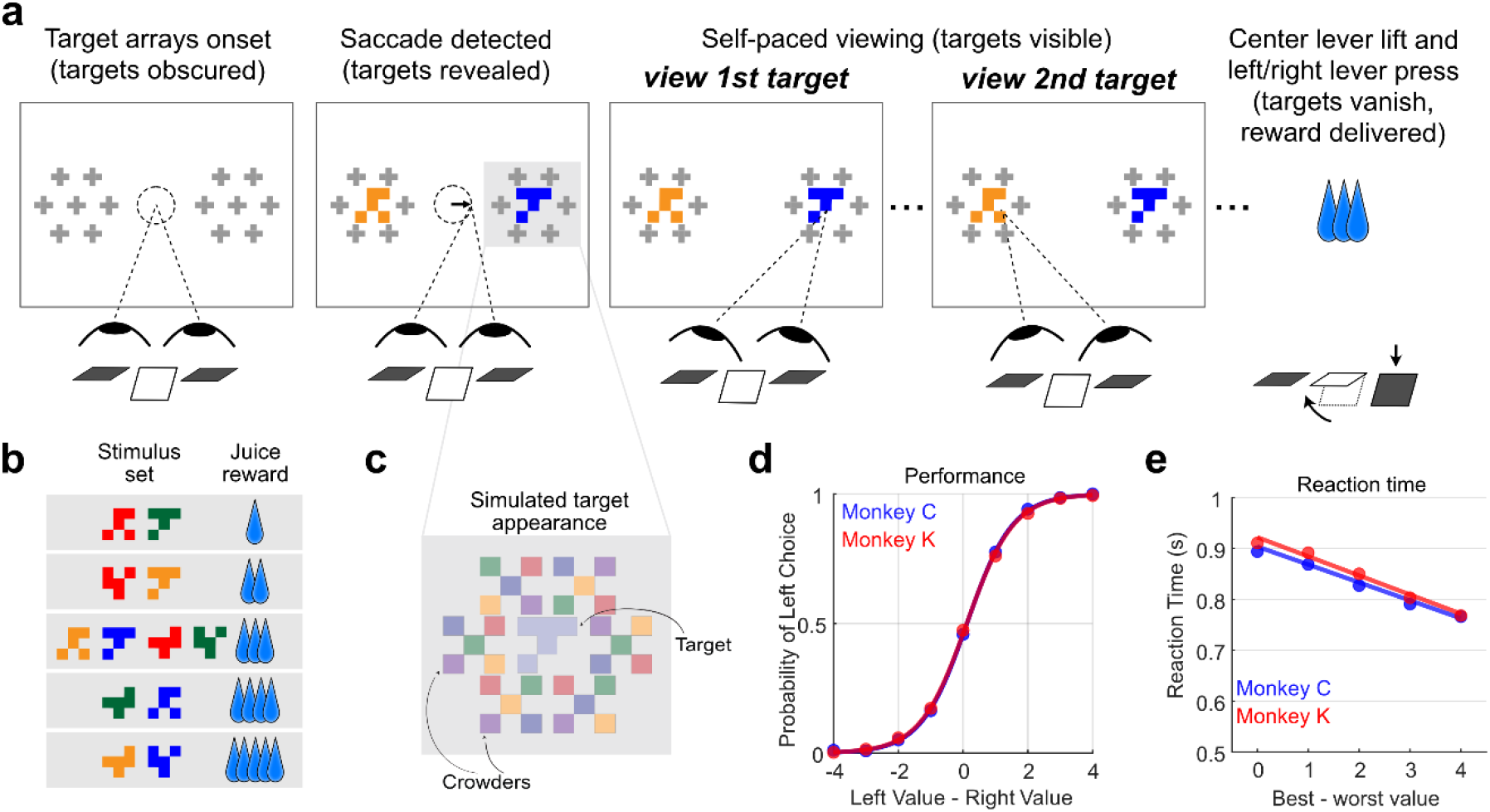
Economic decision-making task and behavioral performance. **(a)** Abbreviated task sequence (not shown to scale), in which monkeys initially fixated on a central dot and then freely viewed the decision targets (the yellow and blue glyphs) before choosing one target by pressing a lever. The ‘+’ shapes indicate the position of visual crowders around the targets; the actual crowders are shown in panel (c). **(b)** Example set of decision targets used in a typical recording session; the associated juice reward for each target (number of drops) is shown on the right. **(c)** Close-up view of an example target (blue, center) surrounded by 6 multi-colored crowders, approximating their appearance on the task display. **(d)** Choice performance: proportion of leti choices as a function value difference between leti and right targets (n=12379 and n=15839 trials for monkey C and K, respectively). Lines show a logistic fit. **(e)** Mean reaction time for each monkey as a function of trial difficulty, defined in terms of the absolute difference in target values; lines show a linear fit.

#### Target properties, visual crowders, and initial mask

The choice targets were colored glyphs, ∼0.75×0.75 degrees of visual angle in size. An example set with associated rewards are shown in Fig. 1b. New target sets with different color and shape combinations were generated every 1-5 sessions to minimize over-learning of the stimuli. Each target was associated with a juice reward ranging from 1-5 drops, with at least two unique targets representing identical reward volumes at each reward stratum (the five horizontal boxes in Fig. 1b). Note that the term ‘target value’ refers to the juice volume associated with a given target, whereas ‘target identity’ refers to a target’s unique appearance (color/shape conjunction).

Two methods were used to encourage the monkeys to fixate directly on both targets before making a choice. First, each target was surrounded by six ‘crowder’ stimuli placed in close proximity, as shown in Fig. 1c. The crowders consisted of multi-colored ‘x’ or ‘+’ shapes that had no association with reward and that were randomly generated for each target array. Crowders reduce the effectiveness of peripheral vision (Crowder & Olson, 2015; Whitney & Levi, 2011), which encourages the monkeys to identify the targets using high-acuity foveal vision – i.e., by shifting their gaze directly onto the targets. Second, the task display was programmed to initially obscure the targets (with a randomly-chosen crowder stimulus) until the monkey initiated their first eye movement outside of the centrally-located initial fixation window. This ensured that the monkey remained unaware of the target values and identities until they initiated a saccade. Once the initial masks were removed, the targets were visible and remained on the screen until the monkey initiated a choice by lifting its hand from the center lever, at which point all targets and crowders were extinguished.

The nominal patch luminance for all the colors used for targets and crowders in Monkey C’s sessions was ∼2.7 cd/m2 with negligible background luminance; and was ∼22cd/m2 for Monkey K’s sessions with a background luminance of ∼4cd/m2, as measured by a Tektronix luminance J17 photometer with J1820 head. Note that during data collection, the luminance of the target stimuli was reduced by up ∼50% of the nominal values, with no change in the crowders, to better obscure the targets (Fig. 1c). This reduction in the luminance did not prevent monkeys from identifying the targets because the choice performance was nearly perfect for the easiest trials (Fig. 1d).

### Neurophysiological Recordings

Linear recording arrays (Plexon V-probes) were introduced into the brain through a sharpened guide tube whose tip was inserted 1-3 mm below the dura. Probes had 16, 24, or 32 channels spaced either 50 or 100μm apart. Depending on the number of probes used, between 32-80 channels were recorded per session (mean 50.0 and 55.6 channels for monkeys C and K, respectively). OFC was identified based on gray/white matter transitions and by consulting a high-resolution MRI acquired from each animal. We targeted the fundus and lateral bank of the medial orbital sulcus and the laterally adjacent gyrus, with anterior-posterior coordinates ranging from +32 to +38 mm anterior to the inter-aural landmark (Öngür & Price, 2000; Saleem & Logothetis, 2012), corresponding approximately to Walker’s areas 11 and 13. Arrays were placed to maximize the number of channels in gray matter.

#### Spike and LFP processing

For all the neural analyses in this study, except for the spectrogram shown in Fig. 2, we defined two types of neural signals for every channel: multiunit activity (MUA) and the high-gamma (HG) signal.

**Fig. 2.**
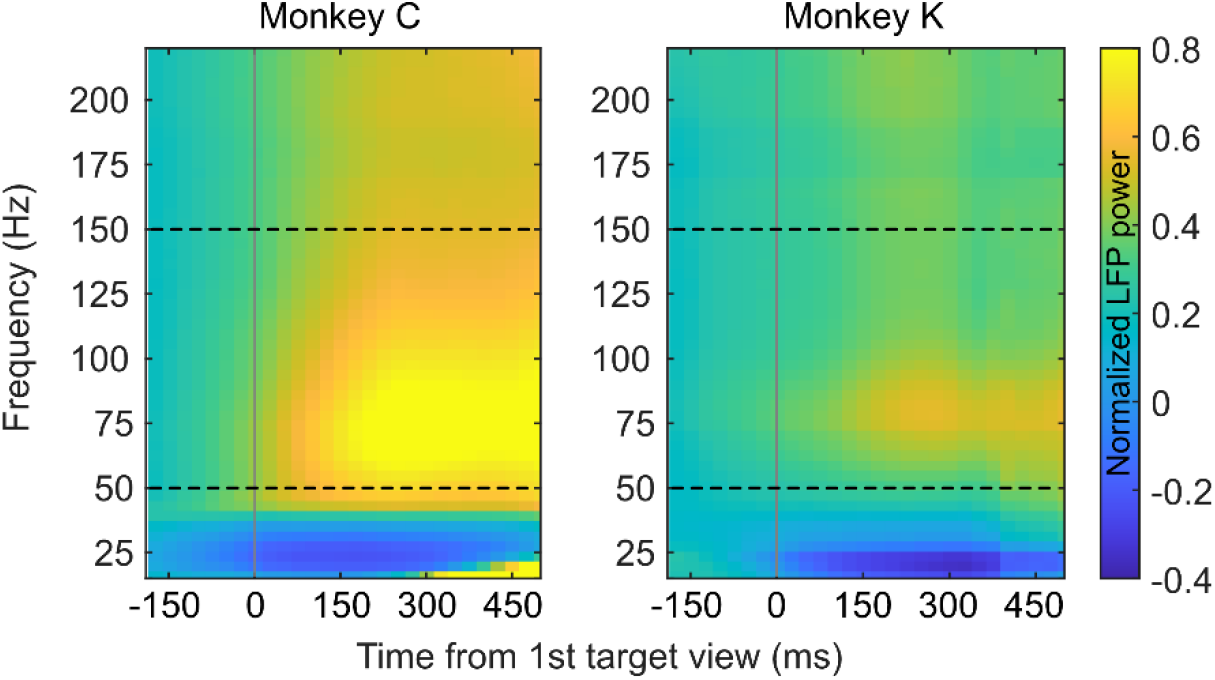
Normalized power spectrogram combined across sessions for each monkey. The power spectrograms are averaged across channels (n= 800 for monkey C, n=1223 for monkey K), aligned to the time at which the first target was viewed in each trial (x-axis). Spectrograms were generated at a single trial level trial using time bins of 500 ms incremented at 100 ms and normalized relative to the baseline (−500 to 0 ms from target onset) before taking the average across trials for each recorded channel in each animal. The heat map color gives the normalized power. Doted lines indicate the 50-150Hz frequency band defining HG in this study.

Spike times were detected online by band-pass filtering the raw signal between 300-1000 Hz, and detecting negative deflections exceeding a threshold of 4 standard deviations below the mean signal amplitude. The MUA signal at each channel is, therefore, unsorted spiking, which reflects the collective activity of multiple nearby neurons. Using unsorted MUA, instead of sorted single-unit activity, permits spiking and HG to be directly compared on a channel-by-channel basis. This is particularly important for interpreting analyses in which HG shows different properties from MUA; if only well-isolated single units were used, then any discrepancy between spiking and HG could potentially be attributed to the influence of unsorted (and therefore unobserved) spike activity. Spike times were counted in 200ms bins, time-locked to the viewing of the decision targets in each trial.

HG signals were obtained by first downsampling the wideband signals from 40 kHz to 1 kHz using an 8th-order Chebyshev Type-I low-pass filter with a cutoff frequency of 400 Hz. The downsampled signal was then notch-filtered at 60, 120, and 180 Hz to remove noise attributable to alternating current power sources. To isolate the high-gamma signal, we bandpass-filtered the notch-filtered data between 50-150Hz using a 1000 order bidirectional FIR filter (MATLAB functions *designfilt.m* and *filtfilt.m*), and the Hilbert transform was applied to the band-passed signal to obtain the instantaneous analytic amplitudes, which we refer to as high-gamma (HG). The 50-150Hz band was chosen because frequencies within this band increase in amplitude relative to baseline after target viewing (Fig. 2) and are comparable to the range for high-gamma used in previous studies (Ray et al., 2008; Rich & Wallis, 2017). Data were arranged into 200ms bins time-locked to the viewing of the decision targets, separately at each bin and for each channel.

To generate the spectrogram shown in Fig. 2, we obtained the full-band LFP from the downsampled signal by applying finite impulse response (FIR) high-pass and low-pass filters of order 1000 at 3 Hz and 300 Hz, respectively. The LFP spectrograms were generated at a trial level with Chronux toolbox in MATLAB (Mitra & Bokil, 2008), using three tapers and a time-bandwidth product of 2. The spectrogram for each trial was then normalized within each frequency with respect to the trial’s baseline, defined as −500 to 0 ms before target array onset. We then averaged the normalized spectrogram across trials to obtain the average spectrogram for each session and channel. Fig. 2 shows the mean spectrogram averaged across channels for each monkey.

### Data Analysis

#### Behavior, task events, and key task variables

We quantified choice performance by a logistic regression in which the fraction of left choices was explained by the value difference between left and right targets (Fig. 1d). Decision reaction times were quantified as a function of choice difficulty, defined as the absolute difference in value between the two targets, with a difference of 0 (equal-value targets) representing the greatest difficulty, and 4, the lowest (Fig. 1e).

In this task, monkeys viewed the targets in each trial sequentially using self-paced saccadic eye movements. The viewing order, total number of target views, and total viewing times were determined entirely by the monkeys. For a detailed description of viewing and choice behavior, see Lupkin & McGinty, (2023).

Analyses were organized with reference to the viewing order of the targets in each trial. Accordingly, the value associated with the first-viewed target is referred to as ‘*value1*’, and the value of the second-viewed target is ‘*value2*’. So that both variables would be defined in every trial, we only used the trials in which monkeys viewed both targets at least once in a trial (96.4% and 90.4% of trials for monkey C and K, respectively).

Because the monkeys’ eye movements were unrestricted, the trial duration and event timing varied across trials. Therefore, the data in each trial were aligned to both the first- and second-target viewing times, and all major analyses were performed in reference to these time points. For some analyses we use “early” and “late” analysis epochs, defined as activity 200-400ms following the onset of fixations onto the first- and second-viewed targets in each trial.

### Coefficient of Partial Determination analysis

#### Single-channel encoding of value and other decision variables

To quantify the encoding of task variables in single channels, we fit the following ordinary least squares linear model at every 200ms time bin (Wilkinson notation):

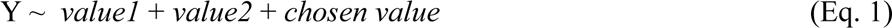

where Y is a trials-by-1 vector of MUA spike counts or HG analytic amplitudes for a given channel and time bin, *value1* and *value2* are the values of the first- and second-viewed targets in each trial (range 1-5 drops), and *chosen value* is the value of the target chosen in each trial (range 1-5 drops). All variables were z-scored across trials before performing the regression. Encoding was quantified by computing for each variable the coefficient of partial determination (CPD) using the methods of Hunt et al., (2018), which captures the unique variance explained by each variable after accounting for the variance explained by all the others. Within each time bin (200 ms, with 40 ms step-size), the mean and SEM of the CPD were calculated across channels, separately for MUA and HG signals; the results are shown in Fig. 4. We use the full regression model in Eq. 1 to calculate the proportion of channels encoding the value variables (Supp. Fig. 2) and to obtain regression estimates used in Fig. 5a, Fig. 6, Fig. 7 & Fig. 10.

**Fig. 3.**
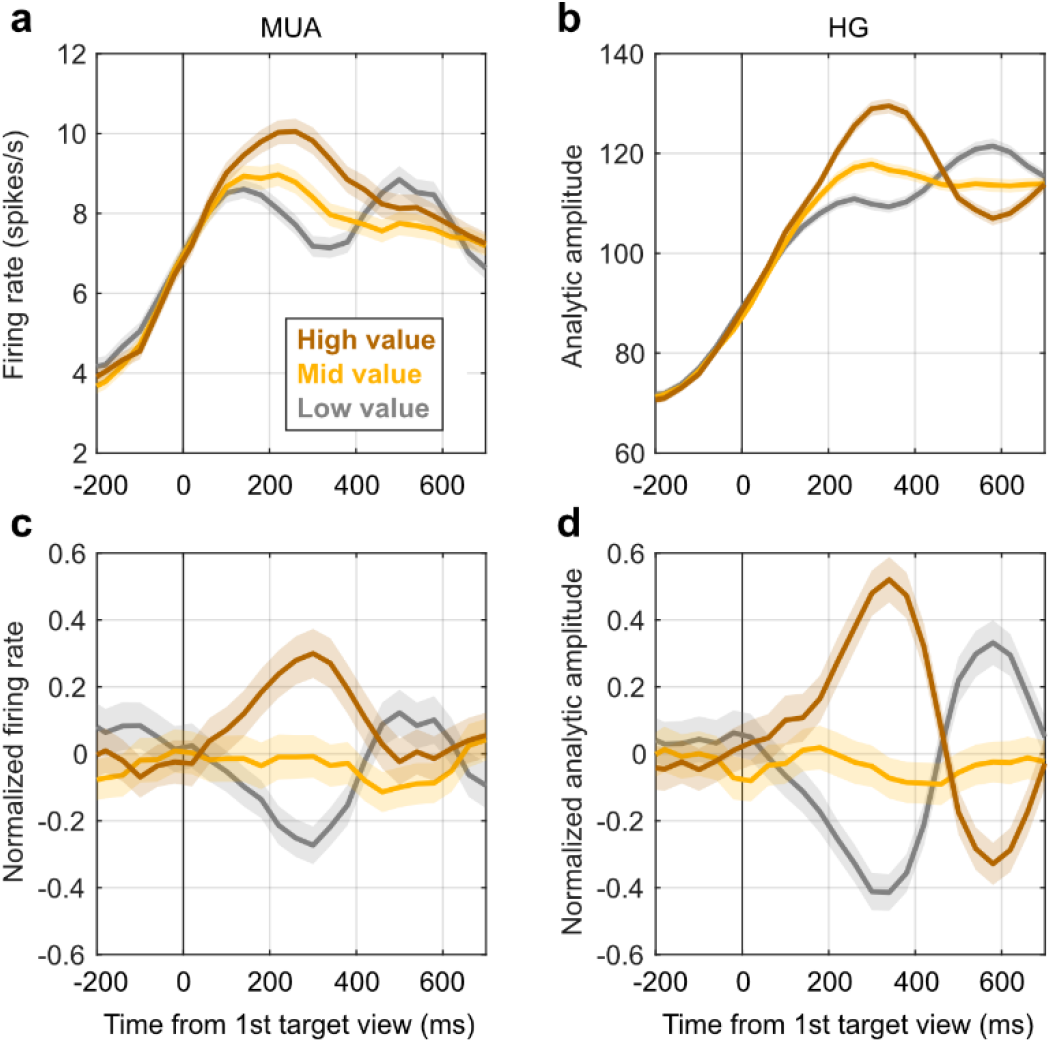
A representative channel showing neural signals modulated by target value. **(a)** Mean MUA firing rate measured on the channel atier viewing the first target, segregated into three trial types: low value (1 or 2 drops of juice), mid value (3 drops) and high value (4 or 5 drops). Shaded regions show SEM across trials (n=233, 235, and 218 for low, mid, and high, respectively). **(b)** Mean analytic amplitude of the HG signal for three trial types. **(c,d)** Same data as (a) and (b) but using neural signals that were z-scored across trials for each time bin.

**Fig. 4.**
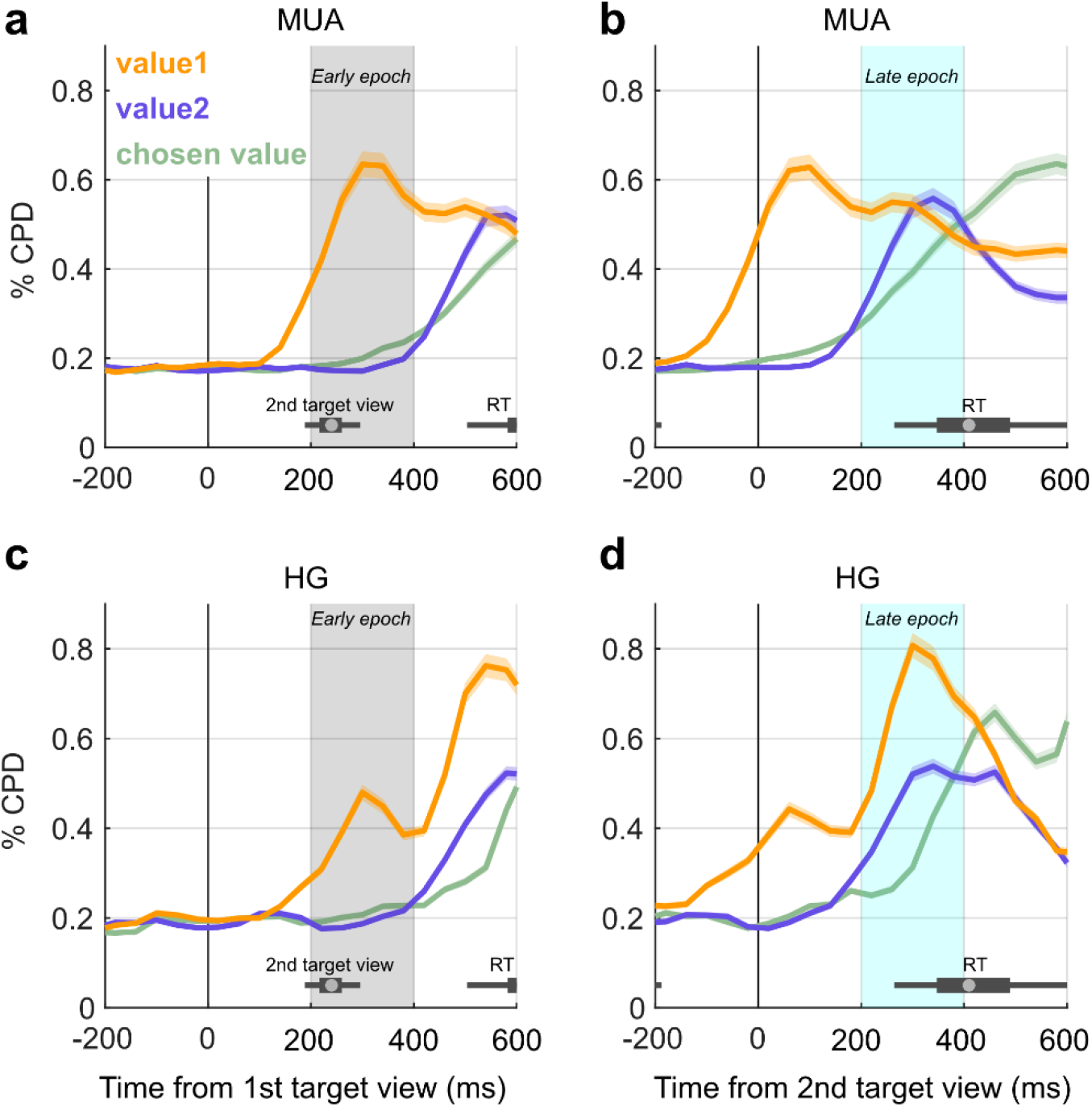
Coefficient of partial determination (CPD) of value variables in MUA and HG. The y-axis in each panel gives the mean CPD of three value variables and the x-axis gives the time relative to viewing the first (a,c) or second (b,d) target. Shading shows SEM across channels (n=2023). Data are combined across monkeys. Panels (a) and (b) show CPD in MUA, and panels (c) and (d) show HG. Shading indicates early (gray) and late (cyan) epochs which are used in subsequent analyses. Horizontal box-plots show the distributions of 2nd target viewing times and reaction times (RT) as indicated. Data separated by monkey are in Supp. Fig. 1, and regression model results showing the percentage of significant channels are in Supp. Fig. 2.

**Fig. 5.**
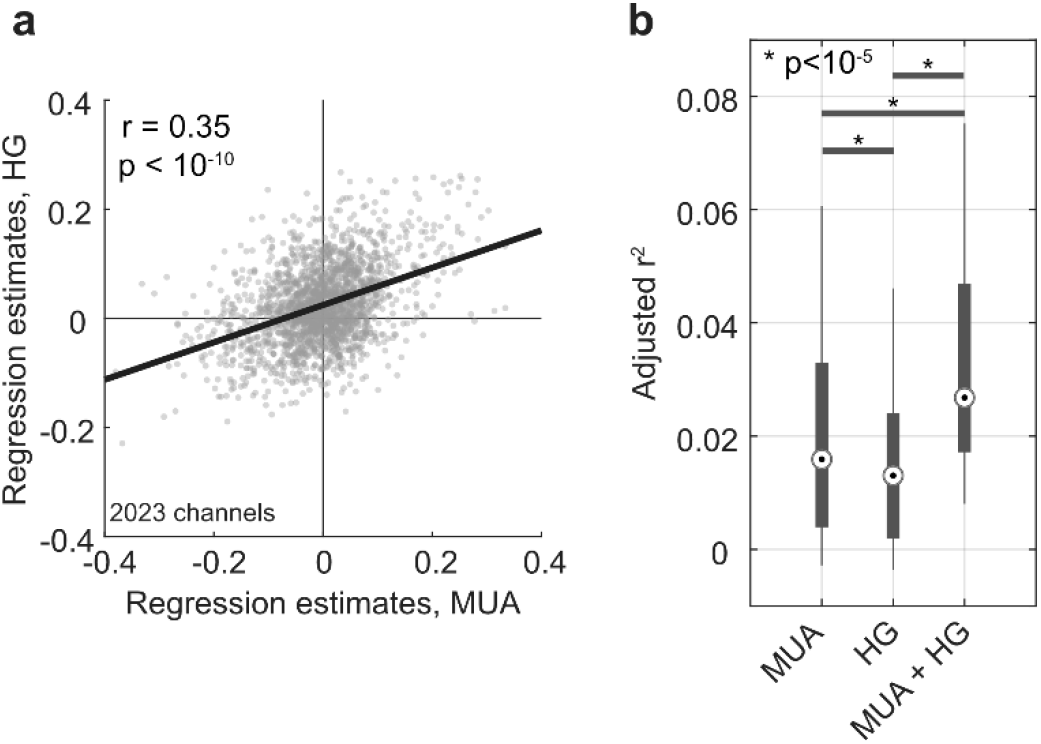
Variance in value explained by MUA and HG. **(a)** Scater plot comparing value encoding in MUA (x-axis) and HG (y-axis) signals. Each dot represents the regression estimates for *value1* for a single channel (Eqn. 1a,b) using data from the shaded epoch in Fig. 4a (n=2023). The black line denotes the least squares regression line. **(b)** Box plots showing the distribution of explained variance in *value1* by MUA, HG, or both signals (see Eq. 2). Data are from the early epoch in Fig. 4a. Only channels with significant effects in either MUA or HG were used (p<0.005 uncorrected in Eq. 1, n = 642); results were similar when using all channels (not illustrated). P-values give the results of a paired t-test across channels.

**Fig. 6.**
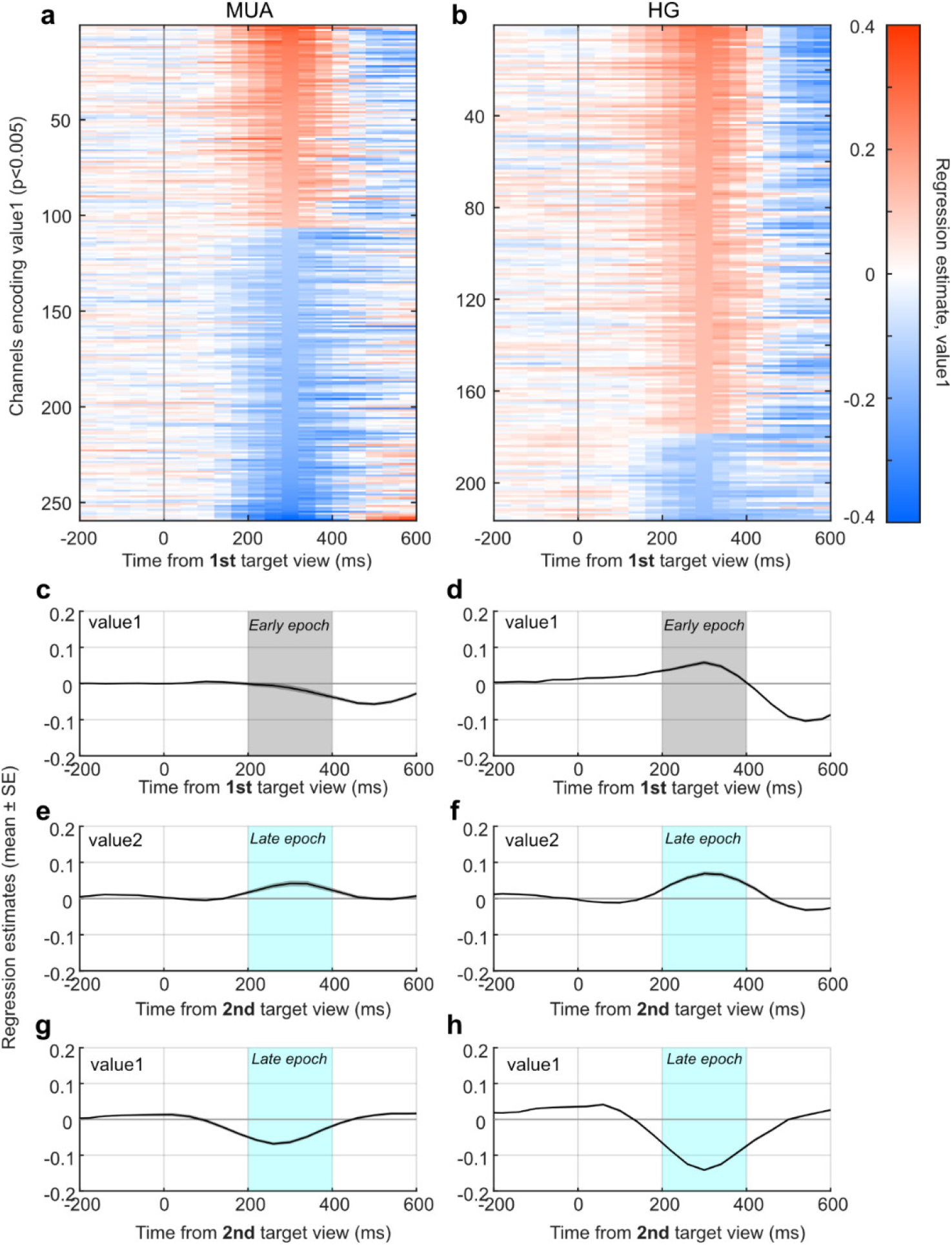
Sign of value coding in MUA and HG. **(a-b)** Regression estimates for *value1* (Eq.1) observed in MUA (**a**) and HG (**b**), ploted in 200ms bins time-locked to the viewing of the first target; only channels with significant value encoding are shown (uncorrected p<0.005, n=259 for MUA and n=216 for HG). Red indicates positive regression estimates (signal increases as a function of value), and blue indicates negative. **(c-h)** Average regression estimates across value-encoding channels. Panels on the leti show average estimates measured in MUA for (c) *value1* atier viewing the first target (n=404), (e) *value2* atier viewing the second target (n=343), and (g) *value1* atier viewing the second target (n=513) Panels on the right (d, f, h) show the same, but for HG signals. Plots using all channels are shown in Supp. Fig. 3. Data from the shaded ‘early’ and ‘late’ epochs were used for analyses reported in the main text.

**Fig. 7.**
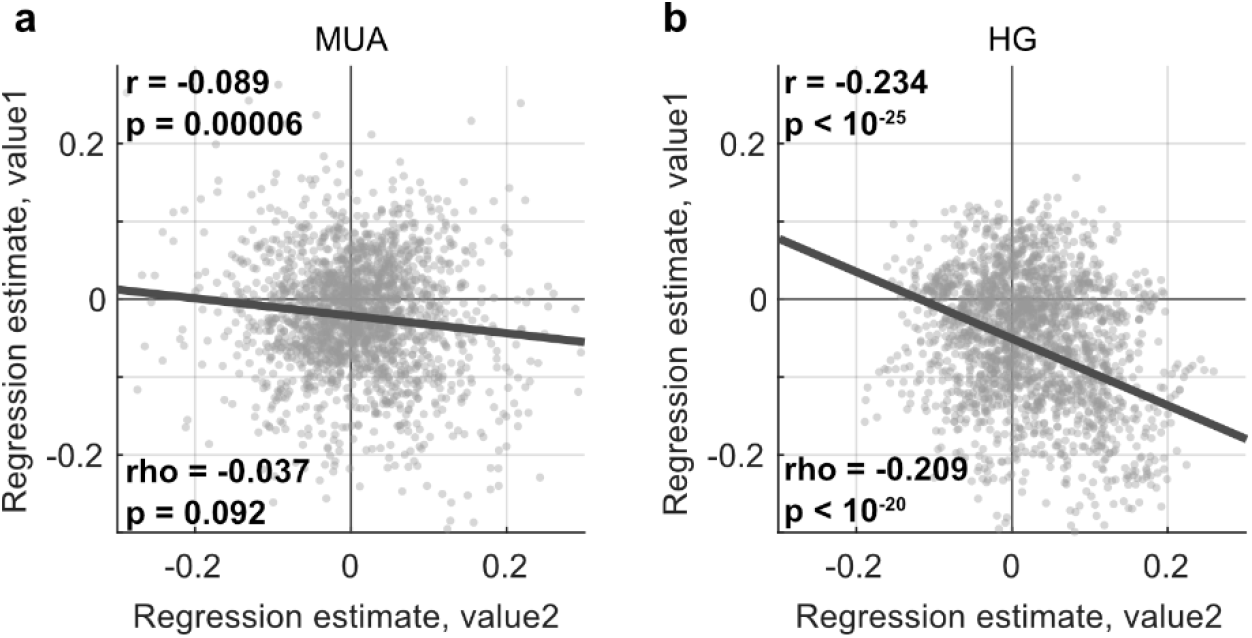
Relationship between simultaneously represented values atier viewing both targets. Channel-by-channel relationship between regression estimates for *value1* and *value2* (Eq. 1) measured atier the second target (late epoch in Fig. 4) in MUA (a) and HG (b). Each dot represents a single channel (n=2023). The ‘r’ and ‘rho’ statistics indicate Pearson’s and Spearman’s coefficients, respectively.

To quantify the unique contribution of MUA and HG to explaining variance in value at the single channel level, we calculated and compared the adjusted r^2^ of three models:

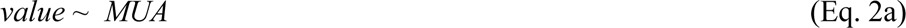

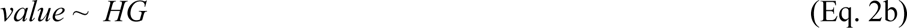

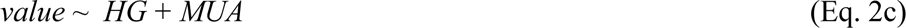

where *value* is either *value1* or *value2*. These regressions were performed at two 200ms time bins: one centered 300ms after the viewing of the first target and the other at 300ms after viewing the second target; these time windows were chosen based upon the time course of value representations observed in Fig. 4 (gray and cyan shaded regions). The difference in adjusted r^2^ between the combined model (Eq. 2c) and the two single-predictor-variable models (Eq. 2a or 2b) was interpreted as the unique variance explained by each variable; adjusted r^2^ is used to account for the extra predictor variable in Eq. 2c.

#### Choice probability analysis in single channels

We used choice probability (CP) analysis to examine the trial-to-trial relationship between neural variability and the variability in choice behavior, an approach first developed for perceptual decisions (Britten et al., 1996) and recently adapted for value-based decision studies (Conen & Padoa-Schioppa, 2015; McGinty & Lupkin, 2023). At an intuitive level, this analysis asks if the activity of value coding signals (MUA or HG) can classify upcoming choices on a trial-to-trial basis while holding all other related variables constant (such as the value or identity of the targets). For in-depth details, see McGinty & Lupkin (2023).

We performed CP analysis separately for MUA and HG at each channel and time bin, separately for each value variable. First, half the trials were used to find the sign of value encoding for *value1* and *value2*, by fitting two one-term linear models. Then, the other half of trials were used for classifying choice outcomes (computing CPs). For each cell and time bin, the CP was computed twice, once using the encoding sign of *value1* to set the positive class, and once using the sign of *value2.* In this way, the CPs are computed with respect to each cell’s encoding of the target values. To compute CPs in the second half of trials, we first normalized the neural signal within each channel and bin by taking the residuals of a two-way factorial ANOVA that explained neural signals as a function of identities of the two targets on each trial. (Identity refers to the 12 unique color/shape combinations.) This procedure removes variance attributable to the target identities and values from the neural signal. The normalized signals were then submitted to a receiver operating characteristic (ROC) analysis, which quantified the ability of an ideal observer to distinguish between trials in which the first or second target was chosen. Classification was only performed over trials in which the target values differed by 0 or 1 because only these trials showed appreciable variability in choice (Fig. 1d).

Classification accuracy was quantified by the area under the ROC curve, where 0.5 indicates chance-level classification. The positive class was set to first-target choices when the value encoding sign was positive, and set to second-target choices when the sign was negative. With this parameterization, an area under the ROC > 0.5 indicates better than chance prediction of choice congruent with a channel’s *value1* sign, and an area < 0.5 indicates better than chance predictions congruent with the *value2* sign (see Fig. 8a and Fig. 8c).

**Fig. 8.**
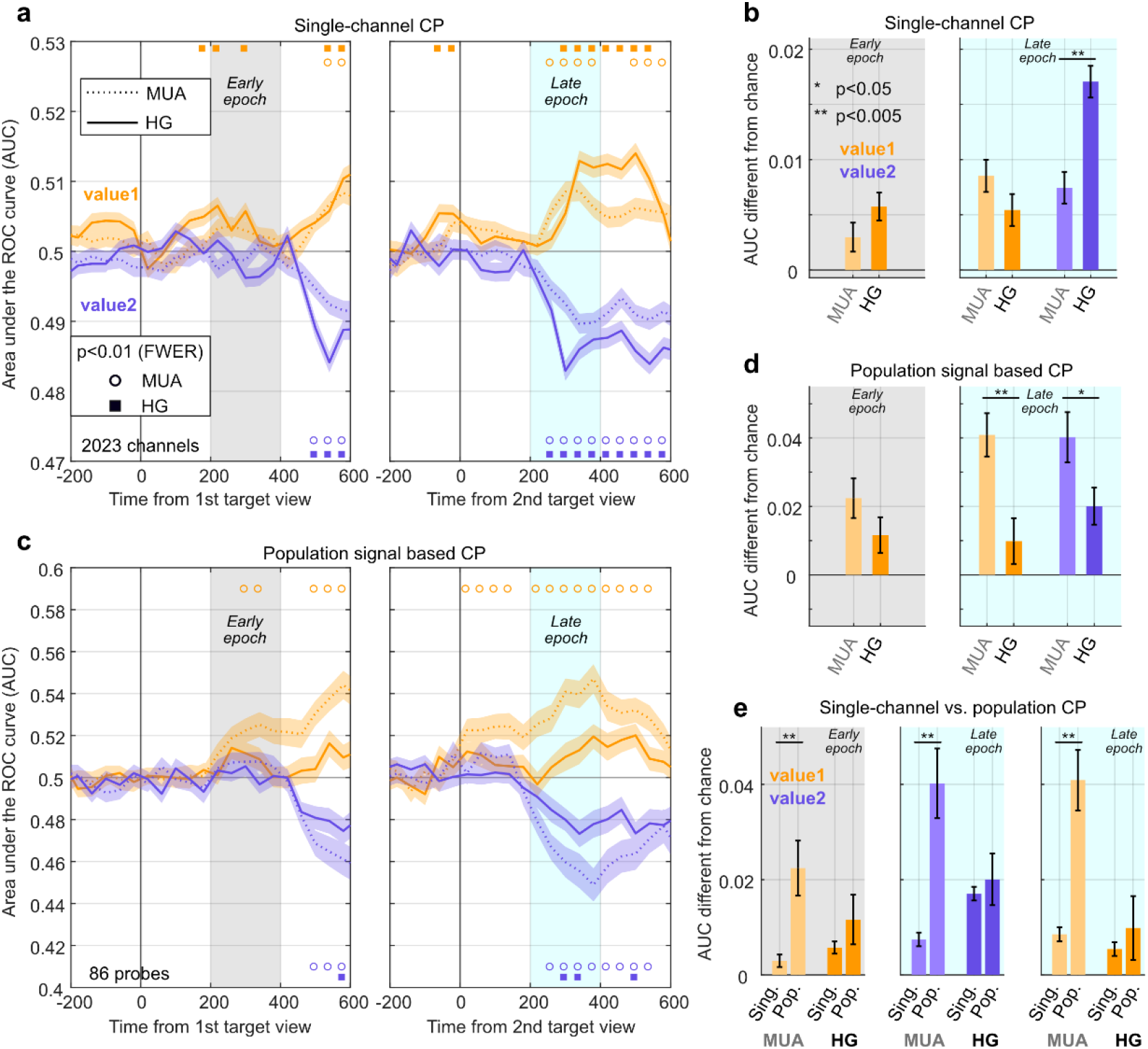
Choice probability analysis for individual channels and population-level value signals. **(a)** Choice predictive accuracy is quantified by the mean area under the ROC curve (AUC), separately for MUA and HG and computed with reference to neural representation of either *value1* or *value2*. Shaded regions indicate SEM across all channels (n=2023) in both monkeys. The x-axes give the time relative to the viewing of the first target (leti panel) and second target (right panel); shaded early and late epochs indicate time windows used for analyses in panels (b), (d), and (e). Shapes at the botom and top of each panel indicate significance (one-sample t-test against chance, p<0.01, FWER corrected across time points), where empty dots indicate significance for MUA and filled squares indicate significance for HG, separately for *value1* (top) and *value2* (botom). In this panel, AUCs represent the accuracy of the ROC analysis for predicting choices in favor of the first target; hence, the AUC for *value1-*based predictions is >0.5, and the AUC for *value2* is <0.5. **(b)** Bar plots showing predictive accuracies relative to chance in MUA and HG in early (leti panel) and late epochs (right panel). In this panel the data for are rectified so that chance performance is 0 and beter-than-chance performance is > 0. Error bars indicate SEM across channels (n=2023). Significance indicators show results of paired t-tests. **(c)** Same as (a) but using population-based value decodes for each probe. Mean, SEMs, and t-tests are calculated across probes (n=86). **(d)** Same as (b) but using data from the population-based analysis in panel (c). Mean, SEM and paired t-tests are calculated across probes. **(e)** The data in (b) and (d) are re-arranged to directly compare single-channel to population-level predictive accuracies for *value1* in the early epoch (leti), *value2* in the late epoch (middle), and *value1* in the late epoch (right). Significance indicator shows results of post-hoc tests from a two-way ANOVA with an interaction term (see main text).

#### Choice probability analysis using population neural signals

We also classified choice outcomes using population-based, single-trial estimates of the target values, obtained by pooling neural activity recorded simultaneously on all channels of a probe. This method used a two-stage, regression-based linear encoder and decoder. In the encoding step, we fit a regression model to explain *value1* or *value2* as a function of neural activity over a set of training trials. In the decoding step, the fitted model is used to estimate *value1* and *value2* using neural activity in held-out test trials. Choice behavior is then predicted in the held-out trials by the submitting neurally-derived value estimates to an ROC classifier. Because this approach requires simultaneously recorded neural signals, these steps were performed using simultaneously recorded signals on all channels of the same probe. In some recording sessions, we recorded multiple probes simultaneously. To address potential violations of statistical independence in averages and t-test results, we confirmed results by randomly selecting one probe per session (see Bootstrap analysis below).

In the encoding step, the data are first stratified into an equal number of training and test trials (chosen at random); the training trials are then used to fit the following L1-regularized linear regression (LASSO) model:

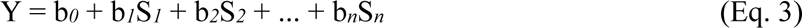

Here Y is either *value1* or *value2*. S*_i_* is the neural signal, (MUA or HG) from channel *i*, b*_i_* is the regression estimate for channel *i*, and n is the total number of simultaneously recorded channels on a probe (16, 24, or 32). The estimates {b*_1_*…b*_n_*} indicate the degree to which the signal of each channel uniquely contributes to explaining the variance in the value variable Y. The model was fit by LASSO regression, which minimizes the mean squared error plus a penalty according to the L1-norm (Hastie et al., 2015).

Before fitting Eq. 3 to *value1* or *value2*, the effects of the non-relevant value variable were removed from the neural data (in only the training trials). This was necessary because the distribution of the *value2* variable was not guaranteed to be identical across every stratum of the *value1* variable; hence, changes in the neural signal attributable to *value2* could be inappropriately attributed to *value1*, leading to inaccurate regression estimates. This step is particularly important for time bins in which more than one value variable is simultaneously represented, (e.g. 200-400ms in Fig. 4b). Therefore, before fitting the model for *value1*, neural data were mean-centered with respect to the identity of the second target; likewise, before fitting the model for *value2*, the neural data were mean-centered according to the identity of the first target.

For decoding, we estimated single-trial population-based value representations by computing the weighted sum of neural signals in the held-out trials:

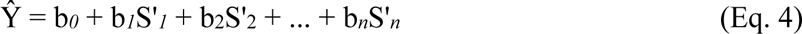

Here Ŷ is the estimated value signal for all held-out trials, b*_i_* indicates the regression estimate from Eq. 3 for channel *i*, b_0_ is the intercept, and S’*_i_* is the neural signal (MUA or HG) from the *i*-th channel in the held-out trials, and *n* is the total number of channels recorded simultaneously on a probe.

We then normalized Ŷ by taking the residuals from two-way factorial ANOVA, in which Ŷ is explained by the identity of the first target and the second target without an interaction term. This removes from Ŷ variance attributable to the trial conditions, leaving only the residual variability in the neurally-derived estimates of *value1* and *value2*. The normalized Ŷ was then used to classify trials according to the decision outcome (first or second target chosen) using a ROC analysis; the normalized Ŷ calculated for held-out trials is referred to as “population value signal” or “population decode” for that trial. ROC classification was performed only over the test trials with target value differences of 0 or 1. The positive class was always defined as choices in favor of the first-viewed target. Therefore, the AUC was above 0.5 when population value signals corresponded to a greater tendency to select the first target, and was below 0.5 when population value signals were associated with a tendency to select the second target.

To compare CPs calculated using single-channel and population value signals, we used 2×2 ANOVA with signal-type (MUA/HG), analysis-type (single-channel/population), and their interaction as factors. Post hoc tests were then performed between relevant pairs using the Tukey-Kramer procedure (MATLAB function *multcompare.m*).

In a follow-up analysis, we compared the variance in value explained by population-level MUA, HG, and combined MUA and HG signals. To do so we computed for every probe the adjusted r^2^ for Eq. 3 when Y was *value1* and S_i_ was MUA on all channels, HG on all channels, or the concatenated MUA and HG on all channels. The time bin used was 200-400ms after second-target viewing. Results were averaged across probes, and are shown in Supp. Fig. 6.

### Correlated variability

We used two approaches to quantify the degree of correlated variability (i.e. shared noise or noise correlation) among simultaneously recorded neural signals. First, we calculated pairwise noise correlations across all channels in a session, using Pearson’s correlations over the baseline signals measured −400 to −200 ms from target onset (i.e. during the initial fixation period). These correlations were performed separately for MUA and HG signals. Second, we used this same data to characterize the structure of shared noise at the level of the population, by performing Principal Component Analysis (PCA) on data from all channels of a probe. When a large portion of noise variance is explained by just a few PCs, this indicates that there are a small number of population-wide activity patterns that dominate the shared noise between channels; in contrast, when more PCs are needed to explain a large portion of variance, it indicates that there are multiple independent activity patterns that each make a small contribution to pairwise noise correlations (Ni et al., 2018). Because the recording probes had different numbers of channels across sessions (16, 24 or 32), we averaged the PCA results within probes of the same type.

### Bootstrap analysis

Because multiple channels were recorded in a session, and because the noise correlations between channels of each probe are positive on average for both MUA and HG, the results from individual channels are not strictly independent, violating the assumptions of many statistical tests. Therefore, to confirm key results we repeated selected analyses using a bootstrap procedure in which we randomly selected one channel at a time from each recording probe (n=86). Results are reported as the median and 95% confidence intervals (CI) of 1000 bootstrap samples. We used a similar procedure to confirm selected CP analysis results, by randomly sampling one probe per session (n=38), and reporting the median and 95% CI over 1000 samples.

## RESULTS

### Task performance

Rhesus monkeys performed a two-alternative, forced-choice decision task in which they chose between two visual targets based on their associated fluid reward (Fig. 1a,b). Visual crowding and a gaze-contingent display (see Methods) were used to prevent the monkeys from discriminating the two targets with peripheral vision, requiring them to fixate directly on the targets to identify them and their associated values in each trial. All neural analyses were therefore performed in reference to the two target viewing times. The choice in each trial was indicated manually by pressing either the left or right response lever located below the task display.

The monkeys looked at both targets at least once in the large majority of trials (96.4% for Monkey C and 90.4% for Monkey K) and their choices and reaction times were consistent with a deliberative process dependent on both target values (Fig. 1d,e). For a detailed analysis of decision and target viewing behavior in this task, see Lupkin & McGinty (2023).

### OFC spiking and high-gamma signals explain unique variance in value

We simultaneously recorded spiking and wideband signals from the OFC during the task. The multiunit spiking activity (MUA) on each channel was obtained from unsorted threshold crossings, and the high-gamma (HG) signal on each channel was defined as the analytical amplitude of the LFP signal between 50 and 150 Hz (see Methods). In this task, the monkeys viewed the two target stimuli in sequence, and all analyses were performed on data time-locked to the viewing of the first and second targets (200ms bins at 40ms increments).

A prior study characterized the value-encoding properties of HG signals in a non-decision, one-option reward expectation task (Rich & Wallis, 2017), but how HG encodes the values of competing value targets during decision-making is unknown. Fig. 3 shows an example of a single channel with activity modulated by the value of the first item the monkey viewed in each trial. In this channel, both the MUA firing rate (Fig. 3a,c) and HG amplitude (Fig. 3b,d) show positive modulation with value around 200-400ms after viewing the target.

To quantify the relationship between the neural signals on each channel and the values of the targets, we fit linear regression models (Eq. 1) at each time bin, separately for MUA and HG. The models explained the neural signals as a function of the values of the two targets in each trial (variables *value1* and *value2*), as well a value variable known to be encoded in OFC spiking activity, *chosen value*, defined as the value of the target chosen in each trial. Encoding for each variable was quantified by the coefficient of partial determination (CPD), which gives the variance explained by that variable after accounting for the effects of all other variables in the model (see Methods).

We found that MUA and HG encoded the value variables with similar time courses. As illustrated in Fig. 4, the variance explained by *value1* increased ∼200ms after the first target was viewed for both MUA and HG (Fig. 4a,c). Likewise, for both MUA and HG, the variance explained by *value2* increased ∼200ms after the second target was viewed (Fig. 4b,d).

However, one notable difference between MUA and HG was the strength of *value1* encoding following the viewing of the second target. To compare encoding across signals and across time points, we defined an “early epoch” and “late epoch”. As illustrated in Fig. 4, the early epoch used activity 200-400ms after the viewing of the first target, and the late epoch used activity 200-400ms after the viewing of the second. For the MUA signal, the explained variance for *value1* peaked in the early epoch (0.64% SEM 0.03, Fig. 3a), and then decreased, becoming significantly lower in the late epoch (Fig. 3b, 0.55% SEM 0.02, p = 0.004 by paired t-test). In contrast, *value1* encoding in HG *increased* in the late epoch, (mean CPD in shaded region of Fig. 3c was 0.48% SEM 0.02, and in shaded region of Fig. 3d was 0.81% SEM 0.03, p<1e-10 by paired t-test.) A similar pattern was evident in the fraction of channels that had statistically significant encoding of these variables (Supp. Fig. 2). Thus, both MUA and HG maintain a significant representation of *value1* after viewing the second target, but only HG showed a significant increase in the explained variance compared to the epoch following the first target.

To compare value encoding in MUA and HG in single channels, we computed the correlation between the estimates from the regression model (Eq. 1). Positive estimates indicate channels for which activity increases as a function of value and negative estimates indicate channels that decrease with value. For *value1*, we compared MUA to HG estimates measured in the early epoch (gray shading in Fig. 4a,c), and also compared MUA to HG estimates in the late epoch (cyan shading in Fig. 4b,d). For *value2*, regression estimates obtained during the late epoch were compared. We found a significant positive correlation between the MUA and HG estimates for *value1* measured on the same channels in the early epoch (r = 0.35, p<1e-10, Fig. 5a). In the late epoch, similar correlations were seen for *value2* (r = 0.39, p<1e-10, not illustrated) and *value1* (r = 0.38, p<1e-10, not illustrated).

These modest positive correlations indicate similar, but not identical modulation by value, suggesting that each signal may explain a unique portion of variance in value relative to the other. We quantified this by computing for each channel the variance in *value1* explained by MUA alone, by HG alone, or by the combination of MUA and HG (Eq. 2), using early epoch activity. The combined signals explained a greater fraction of *value1* variance than either signal alone, as measured by adjusted r^2^ (Fig. 5b). Following the second target (late epoch), a similar result was obtained for explained variance in *value2* and *value1* (not illustrated). Taken together, this indicates that MUA and HG reflect similar encoding of value at each channel, but that each signal also captures a unique portion of the variance in value not explained by the other signal.

### High-gamma increases on average as a function of value

Previously, we have observed that value-coding OFC neurons can either increase or decrease firing as a function of value, with roughly equal fractions of each in recorded samples (McGinty et al., 2016; McGinty & Lupkin, 2023). However, it is unknown whether OFC HG signals have this same property. We quantified the sign of value encoding at each channel using the regression estimates obtained from Eq. 1, where positive estimates indicate channels that increase signal magnitude with value (for example, Fig. 3a between 200-400ms), and negative estimates indicate the opposite.

Consistent with prior studies, the average regression estimates for *value1*-encoding MUA channels were no different from zero after viewing the first target (early epoch, −0.012 SEM 0.008, p=0.13, n=404, for t-test comparison to 0, Fig. 6 a,c). In contrast, for concurrently measured value-encoding HG channels, *value1* estimates were positive on average (0.058 SEM 0.006, p<1e-10, n=404, Fig. 6 b,d), and were significantly greater than the estimates measured in MUA (p<1e-10, n=404, paired t-test). Similarly, *value2* estimates measured after the second target were on average larger for HG than for MUA (late epoch, 0.069 SEM 0.006 vs. 0.042 SEM 0.008, n=343, p=0.0002, Fig. 6 e,f). Finally, for the persistent representation of *value1* after the second target, both MUA and HG showed negative encoding on average (MUA: −0.064 SEM 0.005, p<1e-10; HG: −0.141 SEM 0.003, p<1e-10; Fig. 6 g,h); however, the HG estimates were significantly more negative than MUA estimates (n=513, p<1e-10).

In sum, compared to concurrently observed spiking signals, HG signals were more likely to show net positive or net negative modulation by value, with positive modulation for targets immediately after they are each viewed, and negative modulation for the persistent representation of the value of first target after viewing the second target. We confirmed these findings using all channels (Supp. Fig. 3), and using a bootstrap procedure that ensures independent sampling of activity from each probe (Supp. Fig. 4).

### High-gamma reflects a signature of value comparison

Consistent with prior studies, we found that after the monkeys view both targets, the values of both targets are encoded simultaneously (Fig. 4b,d). We asked whether individual channels tended to encode both variables, and whether for a given channel the encoding of *value1* was related to the encoding of *value2*. We computed correlations between regression coefficients for *value1* and *value2* after viewing the second target (late epoch). For MUA, we found a weak but statistically significant negative correlation between regression estimates (r*_pearson_* = −0.09, p = 0.00006; r*_spearman_* = −0.04, p=0.09; Fig. 7a), consistent with previous reports (Hunt et al., 2018; McGinty & Lupkin, 2023). In contrast, a much larger negative correlation was found for the HG signal (r*_pearson_* = −0.23, p<1e-10; r*_spearman_* = −0.21, p<1e-10; Fig. 7b), indicating that HG channels that were positively modulated by *value2* also tended to show negative modulation by *value1*, and vice versa. We confirmed these findings using a model that uses only *value1* and *value2* as regressors (MUA: r*_pearson_* = −0.09, p = 0.00005; r*_spearman_* = −0.05, p=0.02; HG: r*_pearson_* = −0.30, p<1e-10; r*_spearman_* = −0.29, p<1e-10). In prior studies of single-neuron activity, negative correlations between regression estimates for two target values have been interpreted as a signature of value comparison (see Discussion for details). Thus, this comparison signal is reflected in HG to a much stronger extent than in concurrently observed spiking.

We also compared regression estimates measured after viewing the first target (early epoch in Fig. 4a,c) to those measured after viewing the second target (late epoch in Fig. 4b,d). For both MUA and HG, the early estimates for *value1* were strongly correlated with late estimates for *value2* (MUA: r*_pearson_* = 0.51, p<1e-134; r*_spearman_* = 0.39, p<1e-30; HG: r*_pearson_* = 0.38, p<1e-71; r*_spearman_* = 0.33, p<1e-30; Supp. Fig. 5a), consistent with many channels encoding the currently viewed target with the same sign of modulation. For both MUA and HG the early estimates for *value1* had a small negative correlation with late estimates for *value1* (MUA: r*_pearson_* = −0.10, p<1e-5; r*_spearman_* = −0.02, p=0.36; HG: r*_pearson_* = −0.19, p<1e-16; r*_spearman_* = −0.10, p<1e-5; Supp. Fig. 5b), indicating a slight tendency for channels to switch their encoding sign for *value1* between the early and late epochs.

### Choice-predictive value representations in MUA and HG

In a recent study we showed that variability in value-based decisions could be predicted by population-level representations of value decoded from many simultaneously recorded OFC cells (McGinty & Lupkin, 2023). Here, we use a similar approach to compare the choice-predictive power of MUA and HG value signals.

We first assessed the mean choice-predictive activity of individual channels by computing choice probabilities (CP). For every channel and time bin, we used a set of training trials to find the sign of modulation for encoding *value1*. Then, in the remaining trials, we classified trials according to whether the first or second target was chosen, using an ROC analysis; the positive class was set to first-target choices when the *value1* encoding sign was positive, and set to second-target choices when the sign was negative. In this way, an area under the ROC curve >0.5 reflects classification congruent with the encoding of *value1* for that channel and time bin. The same procedure was repeated using the encoding sign for *value2*; for this analysis, the positive class was set so that an area under the curve of <0.5 reflects classification congruent with the encoding sign for *value2*. Critically, only trials in which the target values differed by 1 or 0 were used for classification because only these trials exhibit significant trial-to-trial variability in choice outcomes (Fig. 1d).

Following the first target, the mean predictive accuracy with respect to *value1* representations was significantly higher than chance level at some time bins for the HG signal, but was consistently near chance for MUA (Fig. 8a, left). After the second target, both MUA and HG showed better than chance classification accuracy (open circles and filled squares in Fig. 8a right). The accuracy for HG was consistently higher than MUA for *value2*-related classification, but not for *value1* (Fig. 8b, right). We confirmed the results using a bootstrap-based sampling method to ensure statistical independence (not illustrated).

We next assessed the predictive accuracy of population-level value signals, which are expected to differ from single-channel activity because they are obtained by taking the weighted sum of activity from all the simultaneously recorded channels in a probe (see Methods and McGinty & Lupkin (2023)). The predictive accuracies for population-level value signals showed the same general temporal pattern as single-channel-based predictions (Fig. 8c), but with greater overall accuracy (Fig. 8e, compare y-axis to Fig. 8a and Fig. 8c). However, population-level value signals derived from MUA were on average more accurate than those derived from HG (Fig. 8d) – the opposite of what was observed in single channels, where HG signals were on average as accurate or more accurate than MUA (Fig. 8b).

Put another way, population-level predictions were more accurate than single-channel-level predictions, but this effect was much greater in MUA than in HG (Fig. 8e). We observed a similar pattern when comparing the average explained variance in *value1* in single channels (Eq. 2) to the variance explained by multiple simultaneous channels (Eq. 3, Supp. Fig. 6). For example, using MUA data from the early epoch, the average explained variance in *value1* for single channels was 2.7% SEM 0.1 (Eq. 2a, results in Fig. 5b), and was much larger for the population-level model (12.1% SEM 1.3, Eq. 3, p<1e-10, by two-sample t-test). In contrast, using the HG data over that same time interval, the mean explained variance for single channels was 1.8% SEM 0.1 (Eq. 2b), but was only slightly more for the population-level model (3.8% SEM 0.5, p<1e-9).

To directly compare population CP with single-channel CP in both signals (Fig. 8e), we used a two-way ANOVA with signal type (MUA/HG), analysis type (single-channel/population), and their interaction as factors. The interaction term was significant for the *value1* results after second-target viewing (late epoch, F_(1,4212)_=7.58, p = 0.006, Fig. 8e, right) and for *value2* results at this same time point (F_(1,4212)_=8.84, p = 0.003, Fig. 8e, middle) indicating that the improved performance of the population decoder compared to single channels was larger in MUA than in HG. For CP effects related to *value1* after the first target (early epoch, Fig. 8e, left) the interaction term was not significant (F_(1,4212)_=2.28, p=0.13), but a post-hoc test revealed a significant difference between single-channel and population-based CP for the MUA data (p=0.012).

Linear decoders like the one used for the population-based model above perform best when the individual signals are independent and contribute unique information about the variable to be decoded (Kohn et al., 2016). The fact that pooling HG signals over many channels results in only a modest increase in decoding performance over single channels suggests that compared to MUA, HG signals measured on different channels of the same probe are more similar (less independent) from trial-to-trial (Hwang & Andersen, 2013).

To test this hypothesis, we measured the noise correlations between pairs of simultaneously recorded channels using activity during the initial fixation period, and compared the correlations for MUA to those for HG. On average, the noise correlations for MUA on pairs of simultaneously recorded channels was 0.072 SEM 0.001 (n=61417 pairs), whereas for HG signals it was 0.326 SEM 0.001 (p<1e-10, paired t-test). We found the same pattern when considering only pairs of adjacent channels (MUA: 0.343 SEM 0.006; HG 0.851 SEM 0.003, n=1988 pairs, p<1e-10) as well as pairs of channels separated by more than 4 array channels (minimum 200µM distance, MUA: 0.050 SEM 0.000; HG 0.270 SEM 0.001, n=53693 pairs, p<1e-10). To characterize the structure of correlated noise across the population, we performed principal component analysis for the baseline activity measured within each probe, and plotted the fraction of noise variance explained as a function of the component number. Compared to MUA, in the HG signal fewer components were necessary to explain a large portion of noise variance (Fig. 9). Taken together, these results show that HG signals have a greater degree of shared covariation across trials, which is dominated by a relatively small number of distinct linear patterns (PCs), compared to concurrently observed MUA.

**Fig. 9.**
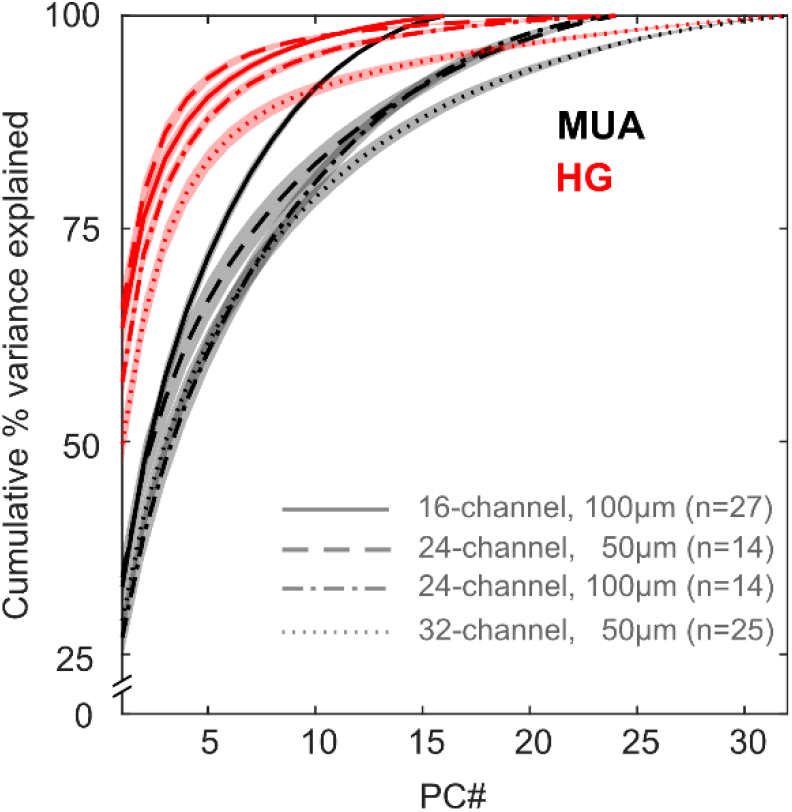
Shared variance across channels in a probe. Cumulative variance explained by principal component (PCs) defined by activity during the fixation period across all channels on a probe. Data are segregated according to probe geometry (number of channels and channel spacing) and neural signal (MUA and HG). Lines and shading show mean and SEM.

### Value and decision signals are accentuated in shallower layers of OFC

Recent studies have reported a higher amplitude of gamma in shallow cortical layers compared to deeper layers in numerous cortical regions in the macaque (Bastos et al., 2018; Johnston et al., 2019; Mendoza-Halliday et al., 2024). Further, another recent study showed that gamma in shallow layers is more dissociable from local spiking activity than gamma in deeper layers (Leszczyński et al., 2020). Therefore, we asked whether the main findings in our study were dependent on cortical depth.

Because the recording probes were typically not inserted normal to the cortical surface, we were unable to estimate the location of specific cortical layers based on electrode location on the probe. Furthermore, methods using electrophysiological signatures (Mendoza-Halliday et al., 2024; van Kerkoerle et al., 2014) were not consistent enough in our hands to confidently assign laminar identities (not shown). Therefore, to obtain a rough proxy for cortical depth, we divided each probe’s channels in half, grouping them into superficial and deep channels. While this coarse division likely misclassifies some channels, the results are consistent with previously identified superficial- and deep-layer properties, as we describe below.

First, we compared the HG amplitude on channels in the two halves of each probe. Consistent with prior reports, the HG amplitude in shallow channels was greater than in deep channels, both after viewing the first and second target (early epoch: 83.4µV SEM 0.7 vs. 66.0µV SEM 0.6, two-sample t-test p=6e-66; late epoch: 81.6µV SEM 0.7 vs. 65.2µV SEM 0.6, p=7e-62) (Bastos et al., 2018; Johnston et al., 2019; Mendoza-Halliday et al., 2024). A similar, but weaker, pattern was observed in MUA in the early epoch (3.62 spikes SEM 0.09 vs. 3.20 spikes SEM 0.10, p=0.002). And the opposite pattern was found in the late epoch (3.16 spikes SEM 0.09 in shallow layers vs. 3.45 spikes SEM 0.09 in deep layers, p=0.019).

We next compared the value encoding properties in shallow vs. deep channels. Shallow channels were more likely to encode value in HG (Eq. 1) than deep channels: For *value1* encoding in the early epoch, 13% of shallow channels encoded value in HG, compared to only 8% of deep channels (chi-square test of proportions, χ^2^=10.92, p=9.5e-4). For *value2* in the late epoch, 13% of shallow and 8% of deep channels encoded value (χ^2^=12.34, p=4.4e-4); and for *value1* in the late epoch coding was significant in 27% of shallow and 15% of deep channels (χ^2^=45.73, p=1.4e-11). Similar trends were observed in MUA, but were not statistically significant for all signals and time points: For *value1* early, encoding was found in 15% and 11% of shallow and deep channels, respectively (χ^2^=8.14, p=0.004). For *value2* late, encoding was found in 11% and 9% (χ^2^=2.2, p=0.13); and for *value1* late, encoding was found in 12% and 10% (χ^2^=1.12, p=0.29) of shallow and deep channels, respectively.

The magnitude of the regression estimates also depended on relative cortical depth. We used a 2×2 ANOVA with an interaction term to compare regression estimates across signals (MUA/HG) and depth (shallow/deep), separately for channels with positive and negative estimates of each value. Among positively-modulated channels, we found a main effect of depth for both *value1* early and *value2* late (*value1*: F_depth(1,2231)_=29.64, p=6e-8; *value2*: F_depth(1,2313)_=17.62, p=3e-5). Post-hoc tests showed that the early *value1* estimates were larger in shallow channels in both MUA and HG (Fig. 10a, left panel, p = 0.0012 and p = 0.0003, respectively). For *value2* in the late epoch, estimates were larger in shallow channels, but only in HG and not in MUA (Fig. 10a, middle, p =0.0004 p = 0.17, respectively). For ANOVAs fit to positive-coding channels, none of the interaction terms were significant (p> 0.23).

**Fig. 10.**
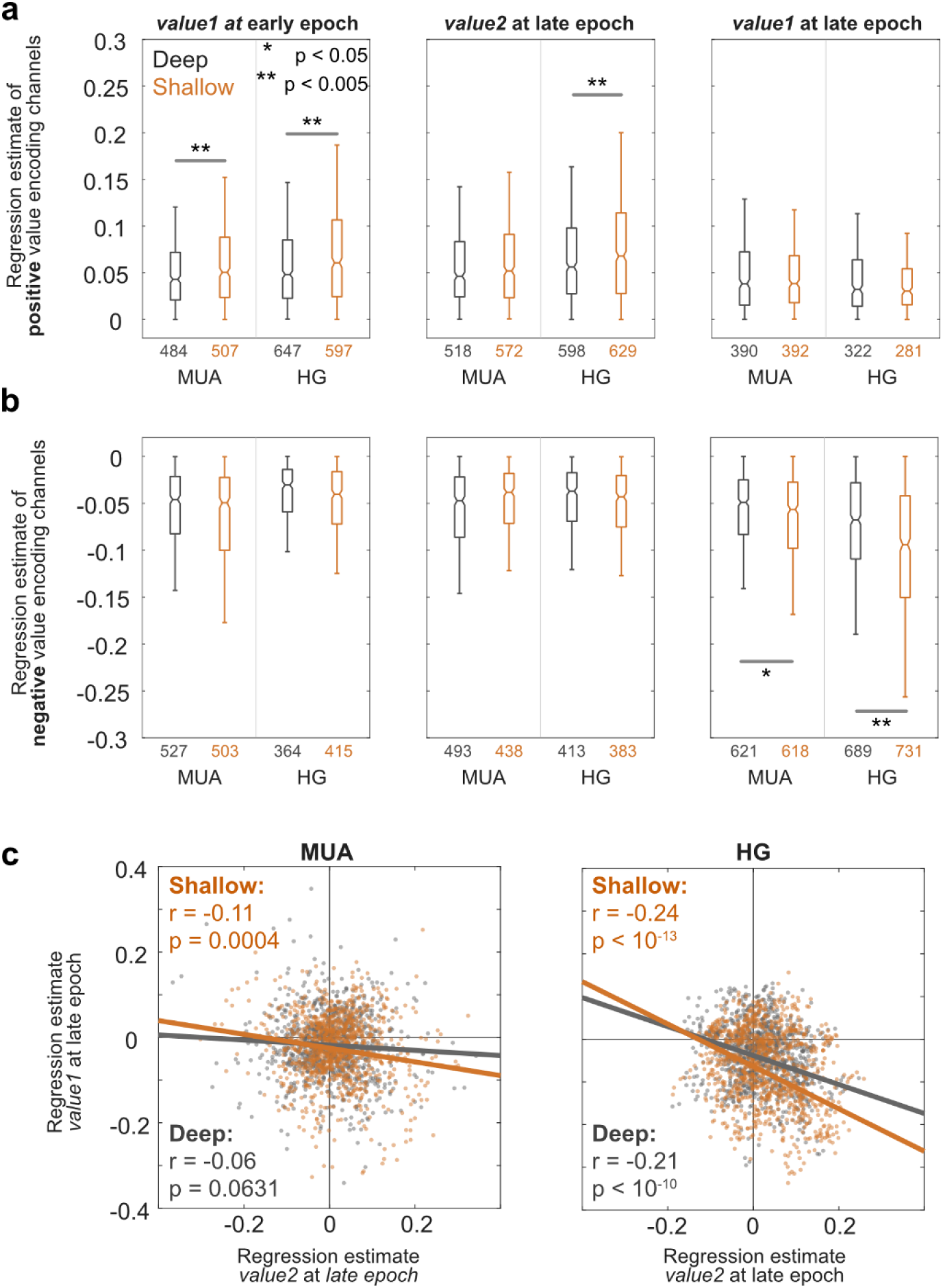
Value encoding properties across relative cortical depth. **(a)** Box plots show regression estimates of channels showing positive value modulation (identified using sign of regresison estimates, Eq.1) for *value1* in early epoch (leti), *value2* in late epoch (middle), and *value1* in late epoch (right), separately for neural signals (MUA and HG) and channel depth (shallow and deep). Early and late epochs indicate 200-400ms time windows atier viewing the first and second target, respectively. The number below each box plot indicates number of channels. Significance indicators show results of ANOVA post-hoc tests. **(b)** Same as (a) but using channels showing negative value modulation. **(c)** Scater plots show relationship between regression estimates of two simultaneously represented values (*value1* and *value2*) atier viewing the second target. Conventions are the same as in Fig. 7, except that data are split by cortical depth.

For negatively-modulated channels, there was no difference in the mean estimates for shallow vs. deep channels for *value1* in the early epoch or for *value2* in the late epoch (Fig. 10b, left and middle panel; p>0.05 for all tests). However, we found a main effect of depth for *value1* in the late epoch (Fig. 10b, right panel; F_depth(1,2655)_=91.4, p=2e-21), as well as a significant interaction term (F_int(1,2655)_=14.8, p = 0.00012). Post hoc tests showed larger negative estimates at shallow vs. deep channels for both HG and MUA (post-hoc test, p=4e-9 for HG and p=0.038 for MUA). However, the significant interaction term indicates that the difference between shallow and deep layers is greater in HG than in MUA – i.e., both signals had larger negative regression estimates in shallow layers, but this effect was accentuated in HG.

Finally, we examined the correlation between *value1* and *value2* regression estimates after the second target (late epoch, as in Fig. 7) as a function of depth, and found a slightly greater negative correlation for shallow channels compared to deeper channels for both MUA and HG (Fig. 10c). However, the difference was not statistically significant as indicated by an *r*-*z*-transform (Z_MUA_ = 1.19, Z_HG_ = 0.60).

## DISCUSSION

Studies in primary sensory cortex show that high-frequency LFPs are in part a function of aggregated spiking activity in local neurons (Ray et al., 2008; Ray & Maunsell, 2011), suggesting that spiking and HG should reflect redundant information about task-related variables or cognitive processes (S. Tremblay et al., 2015). Consistent with this idea, Rich and Wallis (2017) found similar encoding of variables related to reward expectation in spikes and HG signals measured on the same electrodes in monkey OFC. Because primate OFC plays a key role in value-based decisions, we asked whether OFC spiking activity (measured as MUA) and HG would also have similar neural representations for variables specific to decision-making. Although there were many similarities between the two signals, we also identified four major differences in how MUA and HG represented values and decisions.

First, HG signals were more likely to have a consistent sign of modulation by value. Following the viewing of the first decision target, the MUA signals from single channels were equally likely to increase or decrease as a function of the first target value; the same was true for sorted single units in this data set (McGinty & Lupkin, 2023). In contrast, HG signals were much more likely to increase as a function of value. Likewise, following the viewing of the second target, more HG channels increased activity as a function of the second target value, and more HG channels decreased activity as a function of the first target value. The net positive sign for HG value encoding after the first target is consistent with the reward-encoding HG signals observed by Rich and Wallis (2017), as well as with HG signals measured in human OFC (Lopez-Persem et al., 2020). However, Rich and Wallis (2017) also observed a net positive sign in single neuron spiking, and therefore concluded that the two signals exhibited similar modulation by value. In contrast, we observed a clear dissociation, in which HG, but not spiking, consistently increased as a function of values.

Second, in the MUA signal, the variance explained by the first target value reached a maximum ∼300ms after viewing the first target, and then decreased following the second target. By contrast, the HG signal showed the opposite pattern: explained variance increased following the first target, and then increased again and reached a peak following the second target. In other words, for the representation of the first target value the HG signal was dissociated from the spiking signal not just in terms of the sign of modulation (see above), but also in the time course of explained variance.

Third, following the second target, HG signals that were positively modulated by the second target value tended to also be negatively modulated by the first target value, resulting in a negative correlation between regression estimates for the two values. In contrast, only a very weak negative correlation was evident in MUA, consistent with prior reports showing a small or no relationship between regression estimates for serially presented decision targets (Ballesta & Padoa-Schioppa, 2019; Hunt et al., 2018; M. Z. Wang et al., 2022). A negative relationship between regression estimates of simultaneously represented value variables has been interpreted as a form of value comparison, possibly resulting from mutual inhibitory interactions between populations of value-encoding neurons (Strait et al., 2014; Machens et al., 2005; X.-J. Wang, 2002). This interpretation has been questioned on the grounds that negative correlations can be an artifact of the experimental design – specifically to unequal value ranges for the two targets, as explained in Ballesta & Padoa-Schioppa (2019). However, such an artifact does not explain our data: First, the value ranges for both targets were equal (1-5 drops). Second, if the negative correlation were an artifact, it would be equivalent in MUA and HG.

Fourth, HG signals differed in their ability to predict decision outcomes in single trials. As we showed recently, the average predictive accuracy of individual value-coding OFC neurons is near chance, even over a very large sample of cells; however, pooling the activity of simultaneously observed neurons to compute a population-level value signal results in predictions that are far more accurate (McGinty & Lupkin, 2023). In this study, single-channel HG signals could predict choices as well or better than MUA observed on the same channels, consistent with (Hwang & Andersen, 2013), but the opposite was true for population-level value signals computed using simultaneously observed activity. In other words, for MUA, predictive accuracy was significantly improved for population vs. single-channel signals, whereas for HG the improvement for population signals was marginal.

In summary, while OFC spiking and HG signals had many similarities, consistent with prior studies, HG signals also exhibited unique value- and decision-related properties which were not apparent from concurrently measured spiking.

### Potential mechanisms dissociating MUA and HG

The biophysical origins of the spike-HG dissociations we observed are unclear. Consider the observation that HG signals tend to encode the value of the immediately viewed target with a net positive sign, whereas the MUA encoding sign is on average zero. It is sometimes assumed that HG reflects an aggregate of local spiking activity (Ray et al., 2008; Ray & Maunsell, 2011; Rich & Wallis, 2017). However, if this were strictly the case then the HG signals in this study might be expected to be evenly split between positive- and negative-signed channels, or to show no significant encoding due to aggregation of opposing activity patterns.

A likely explanation for this dissociation is that HG also reflects the aggregation of other local neuronal events, such as synchronized subthreshold somatic and dendritic potentials, related to synaptic transmission (Buzsáki et al., 2012; Leszczyński et al., 2020). Consistent with this idea, gamma signals are stronger in the superficial cortical layers than deeper layers (Mendoza-Halliday et al., 2022), and gamma in superficial layers is more dissociated from neuronal spiking than gamma in deeper layers (Leszczyński et al., 2020). We found that HG power and the tendency to encode value were both greater in the shallower half of array channels, and that regression estimates among positive-coding channels were higher on average in the superficial half of the array. If the hypothesis of Leszczyński et al., (2020) is correct, then our results suggest that the spike-dissociable features of value encoding in OFC HG are primarily due to synaptic inputs to superficial layers. To test this hypothesis would require recordings in which cortical layers are fully resolved (unlike the approximate demarcation used here), and in which the inputs to these layers are identified and examined with respect to value-coding properties.

We now consider a second dissociation: HG explained *value1* variance more after viewing the second target compared to after viewing the first target. In our task, animals view targets sequentially. While viewing one target, the other target is obscured due to being distant and due to crowders placed around the cue (Fig. 1). Thus, optimal performance requires that one target value be remembered while viewing the other. In other words, the monkeys must use some form of working memory.

It has been reported in multiple studies that gamma signals play a significant role in working memory maintenance and executive control (Lundqvist et al., 2011, 2016, 2018; Miller et al., 2018); but also see Thrower et al., (2023). These studies find that gamma power in dorsolateral prefrontal cortex increases as a function of memory load, and propose that gamma reflects short-term changes in synaptic weights that encode the memoranda. In our study, gamma-range signals explain the most variance in the first target value at the time point when it must be maintained in memory. This result suggests that decisions between sequentially viewed items may rely on mechanisms similar to those used in classic working memory tasks, with high-gamma LFPs having a similar role. Because our task did not have mandated delay periods, we could not identify processes related to working memory maintenance distinct from those related to the decision. Future decision studies may be able to test this hypothesis with task designs like those used by Miller and colleagues (Miller et al., 2018), using mandated delay periods to assess the dynamics of OFC HG and spiking as value information is held in working memory (Enel et al., 2020).

### Role of shared noise in value coding and choice-predictive properties of HG

A fundamental property of neural networks is the degree of shared noise or “noise correlation”, defined as the tendency for two signals to fluctuate together from moment to moment after accounting for all other observable variables. The magnitude and structure of these correlations can have complex and subtle effects on the information that can be decoded from single neurons and neural populations (Kohn et al., 2016).

Noise correlations between pairs of neurons in primate neocortex are typically in the range of r=0.01 to 0.1 (Cohen & Kohn, 2011), and we recently reported a mean noise correlation of 0.019 in sorted OFC neurons from this same data set (McGinty & Lupkin, 2023), consistent with a prior study by Conen & Padoa-Schioppa (2015). In the present study, the pairwise noise correlations among HG signals were much higher than among MUA signals measured on the same channels (mean of 0.326 vs. 0.072). The shared noise among HG signals was also lower-dimensional than MUA signals (Fig. 9), meaning that for HG an equivalent portion of the noise variance could be explained by a smaller number of linear patterns of activity fluctuation. These differences are consistent with the fact that HG signals are inherently less “local” than MUA signals, i.e. that they appear reflect activity over a larger spatial extent (Dubey & Ray, 2016; Kajikawa & Schroeder, 2011; Katzner et al., 2009; Xing et al., 2009).

The higher noise correlations in HG compared to MUA help explain the differences in the choice-predictive properties for the two signals. According to Haefner et al (2013), for a sufficiently large pool of neurons with net positive shared noise, the CP for a given neuron mostly reflects its correlation with the other neurons in the pool rather than that neuron’s individual contribution to the choice; the greater the shared noise, the higher the noise correlation for single cells. This is consistent with our observation that single-channel HG signals had CPs that were equal to or greater than the concurrently measured MUA.

The high shared noise in HG signals may also explain the somewhat paradoxical finding that pooling HG signals does not substantially increase the accuracy of CPs or the explained variance in the value signal (unlike for single-cell spiking or MUA). Because HG channels are more correlated (i.e. less independent), the information gained by pooling two or more HG channels is small compared to the information gained by pooling MUA channels. One limitation to this conclusion is the linear geometry of the vertical recording arrays, which placed the channels within a relatively small volume – e.g. 32-channels within a 1.6mm track. A more spatially-distributed set of channels would be expected to have overall lower shared noise, and as a consequence may improve the performance of pooled-signal decoders like the one used here.

### Implications for decision mechanisms and decision BMIs

Our results suggest that HG provides access to features of OFC value signals that would be difficult to detect in the activity of single neurons. Because OFC neurons have a heterogeneous value code (are equally likely to increase or decrease firing as a function of value), representations of low and high value targets are expected to be similar in terms of the overall amount of neural activity evoked across the OFC, which could limit the information that could be decoded without accessing spike-level activity. However, our results show that HG signals are much more uniform, increasing on average as a function of value. Likewise, compared to MUA, HG signals showed a larger negative correlation between regression estimates for the two values, which is interpreted as a form of value comparison that is critical for binary choices (Hunt et al., 2018; Strait et al., 2014).

The fact that these two critical signals can be accessed without measuring single-neuron activity has implications for the development of a brain-machine interface (BMI) for decision-making and other forms of motivated behavior that depend on OFC. The development of BMIs for sensory and motor cortex have often leveraged neural signals decoded from pooled spiking activity across array channels (Andersen et al., 2022). In contrast, our present findings suggest that in OFC, important signals may not be easily accessible from spiking, but are better accessible in HG. HG signals from implanted arrays are less susceptible to decreases in signal quality that typically occur over time (Stavisky et al., 2015). Furthermore, we found that HG value signals were stronger in superficially-located recording channels, consistent with recent studies of HG and MUA properties in different layers (Leszczyński et al., 2020). Together, this suggests that value and decision-related signals could be obtained with less invasive approaches and could be more robust over the lifetime of a BMI implant. Important directions for future work are the nature of HG signals that can be measured in behaviorally more complex decision settings, as well as the potential spike-HG dissociations in other prefrontal areas that have a high degree of heterogeneous encoding.

## Acknowledgements

W.T. Newsome for funding and material support; J. Brown, E. Carson, S. Fong, A. McCormick, M. Ortiz, J. Powell, J. Sanders, D. Siegel for technical assistance; and D. Headley and B. Krekelberg for helpful comments on the manuscript.

## Funding

This work was supported by the Howard Hughes Medical Institute (W.T. Newsome), National Institutes of Health Grant K01-DA-036659-01 (V.B.M.), the Busch Biomedical Foundation (V.B.M.), and a Whitehall Foundation fellowship (V.B.M.).

## Author contributions

DS and VBM conceived the study. VBM designed the experiment. SML and VBM collected the data. DS analyzed the data. DS and VBM drafted the paper. SML edited the draft. DS and VBM wrote and edited the final draft.

## SUPPLEMENTARY FIGURES

**Supp. Fig. 1.**
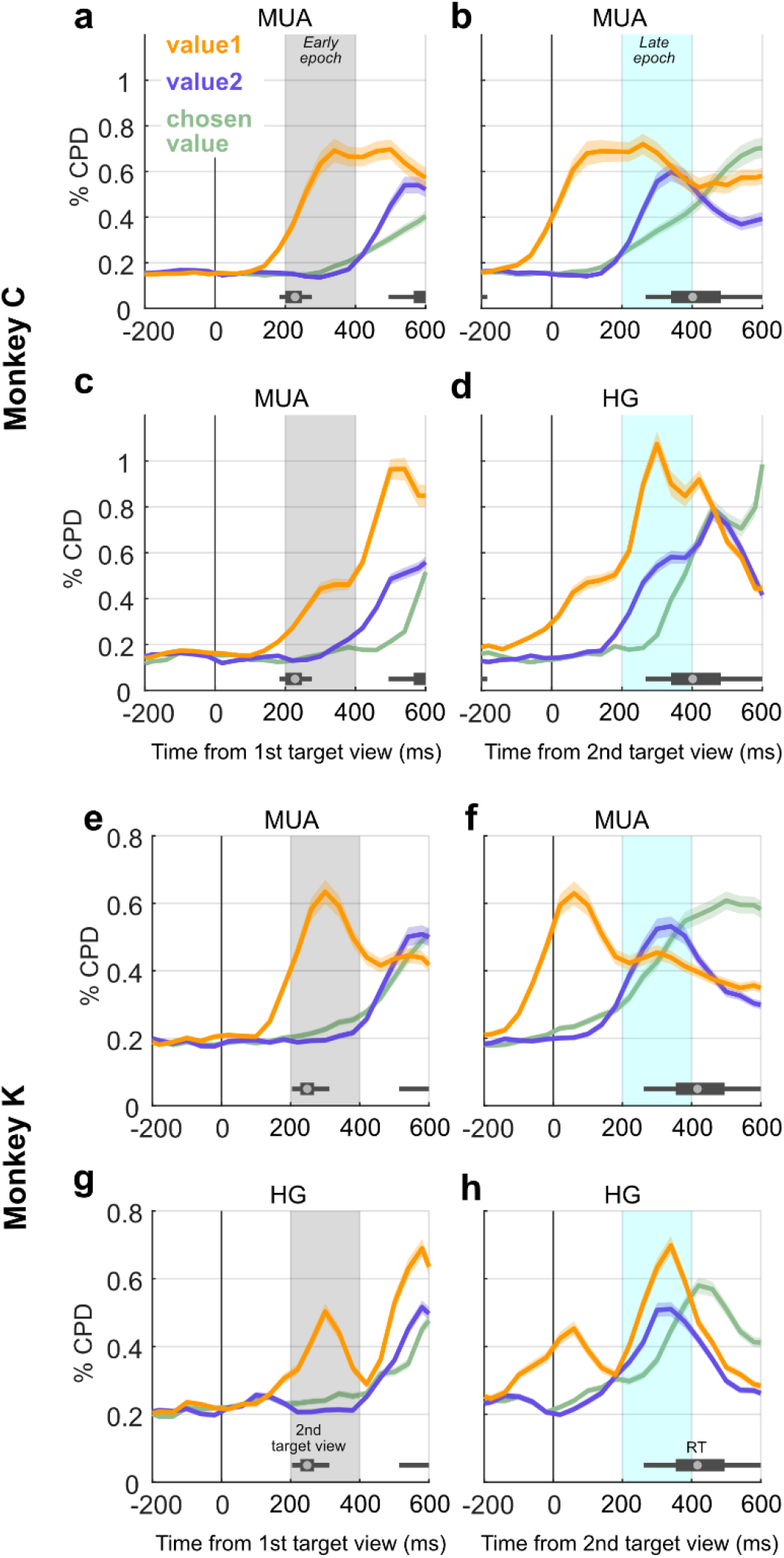
Coefficient of partial determination (CPD) of value variables in MUA and HG, separately for each monkey. Conventions are the same as Fig. 4, but data are shown separately for Monkey C (a-d) and Monkey K (e-h).

**Supp. Fig. 2.**
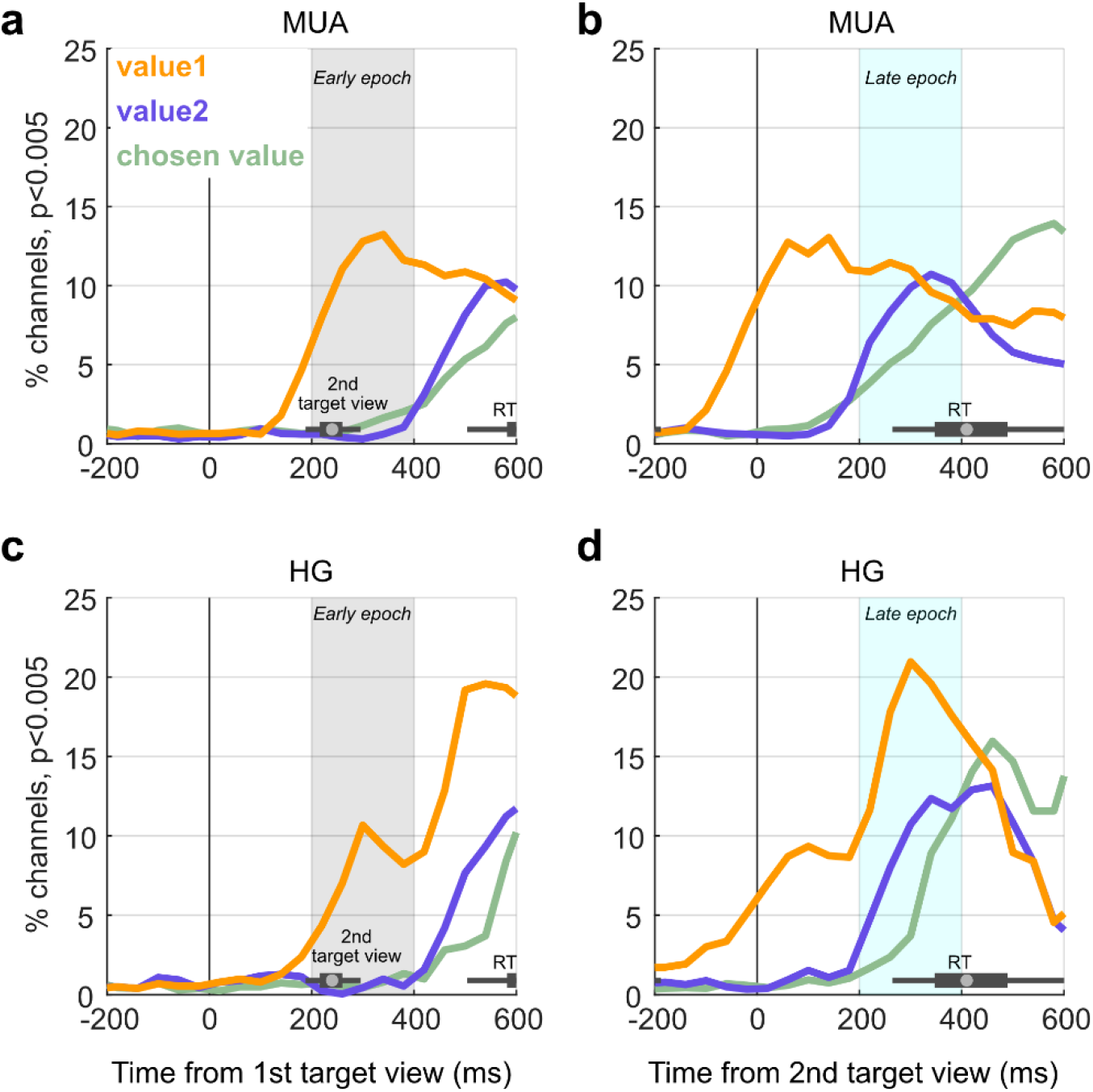
Percent significant channels for value variables in MUA and HG. The y-axis in each panel gives the percentage of channels significantly encoding task variables in the regression shown in Eq. 1 (p<0.005, uncorrected) and the x-axis gives the time aligned to viewing the first (a,c) or second (b,d) target. Data are combined across monkeys. Other than the different data shown on the y-axis, conventions are the same as in Fig. 4.

**Supp. Fig. 3.**
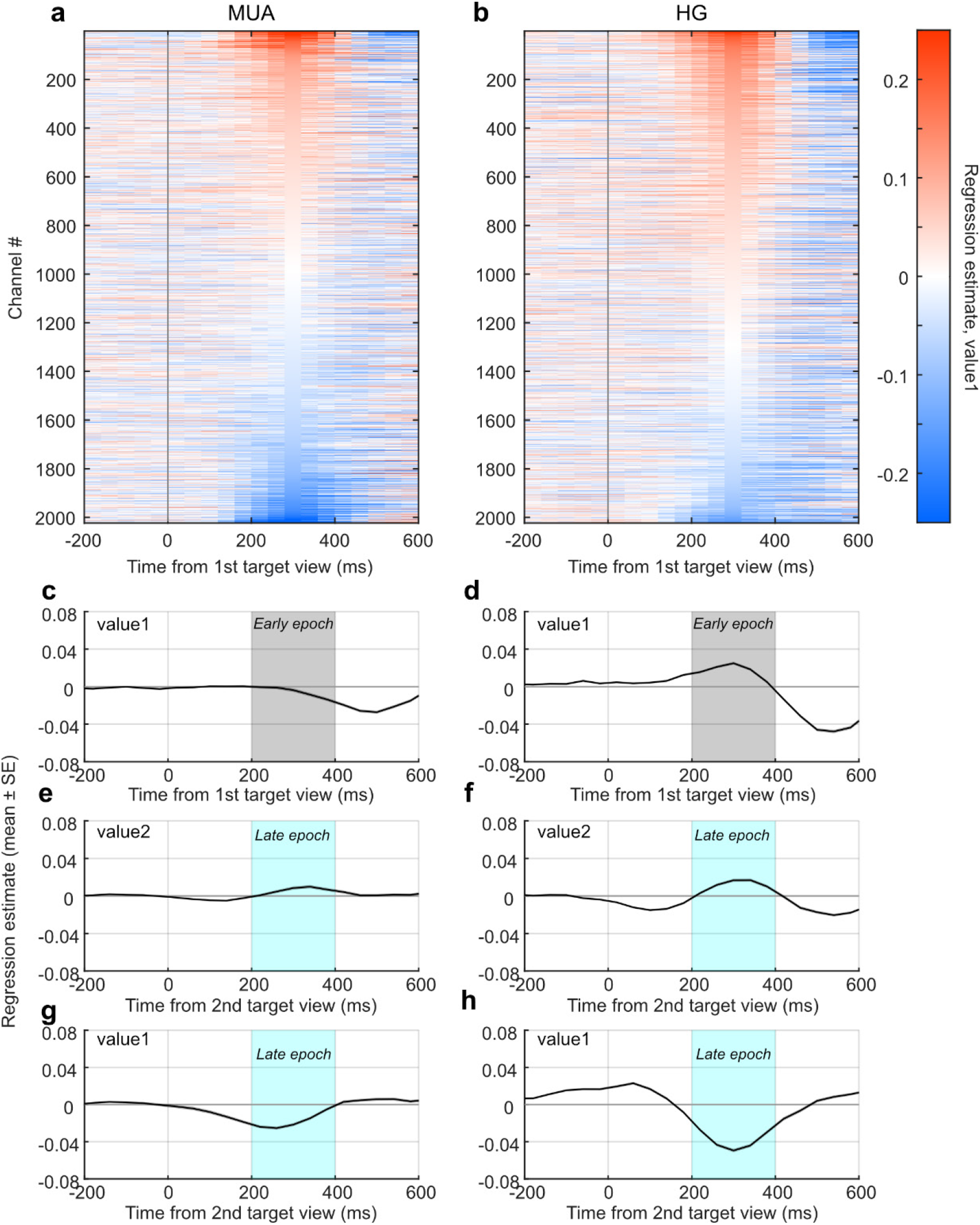
Sign of value coding in MUA and HG using all channels (n=2023). Conventions are the same as Fig. 6. Mean regression estimates for *value1* in the early epoch (panels **c-d**): −0.004 SEM 0.002 for MUA (p=0.052, t-test against 0) and 0.025 SEM 0.002 for HG (p=1.1e-54 against 0) (paired t-test for MUA vs. HG, p=1.5e-49). Similarly, mean regression estimates for *value2* in the late epoch (panels **e-f**): 0.008 SEM 0.002 for MUA (p=2.1p-6) and 0.017 SEM 0.002 for HG (p=7.1e-23) (MUA vs. HG, p=1.1e-5). Mean regression estimates for *value1* in the late epoch (panels **g-h**): −0.022 SEM 0.002 for MUA (p=1.2e-35) and −0.049 SEM 0.002 (p=2.1e-137) (MUA vs. HG, p=2.3e-45).

**Supp. Fig. 4.**
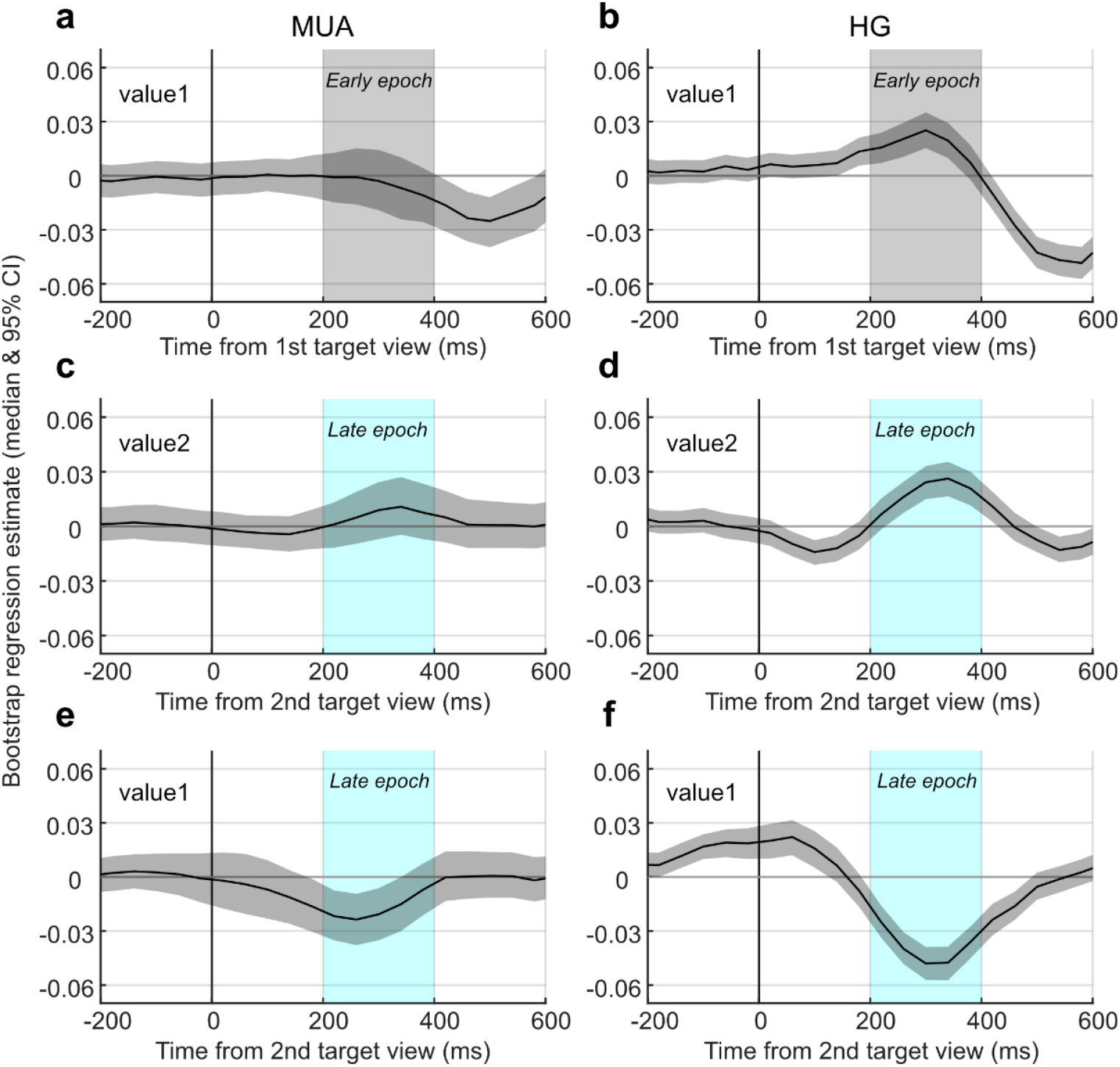
Bootstrap analysis results for the sign of value modulation in MUA and HG, related to Fig. 6. Each panel shows the median and 95% CI of the regression estimates of a single channel taken each probe (n=86) iterated 1000 times. Panels on the leti **(a,c,e)** are for MUA and panels on the right **(b,d,f)** are for HG. Otherwise, conventions are the same as in Fig. 6.

**Supp. Fig. 5.**
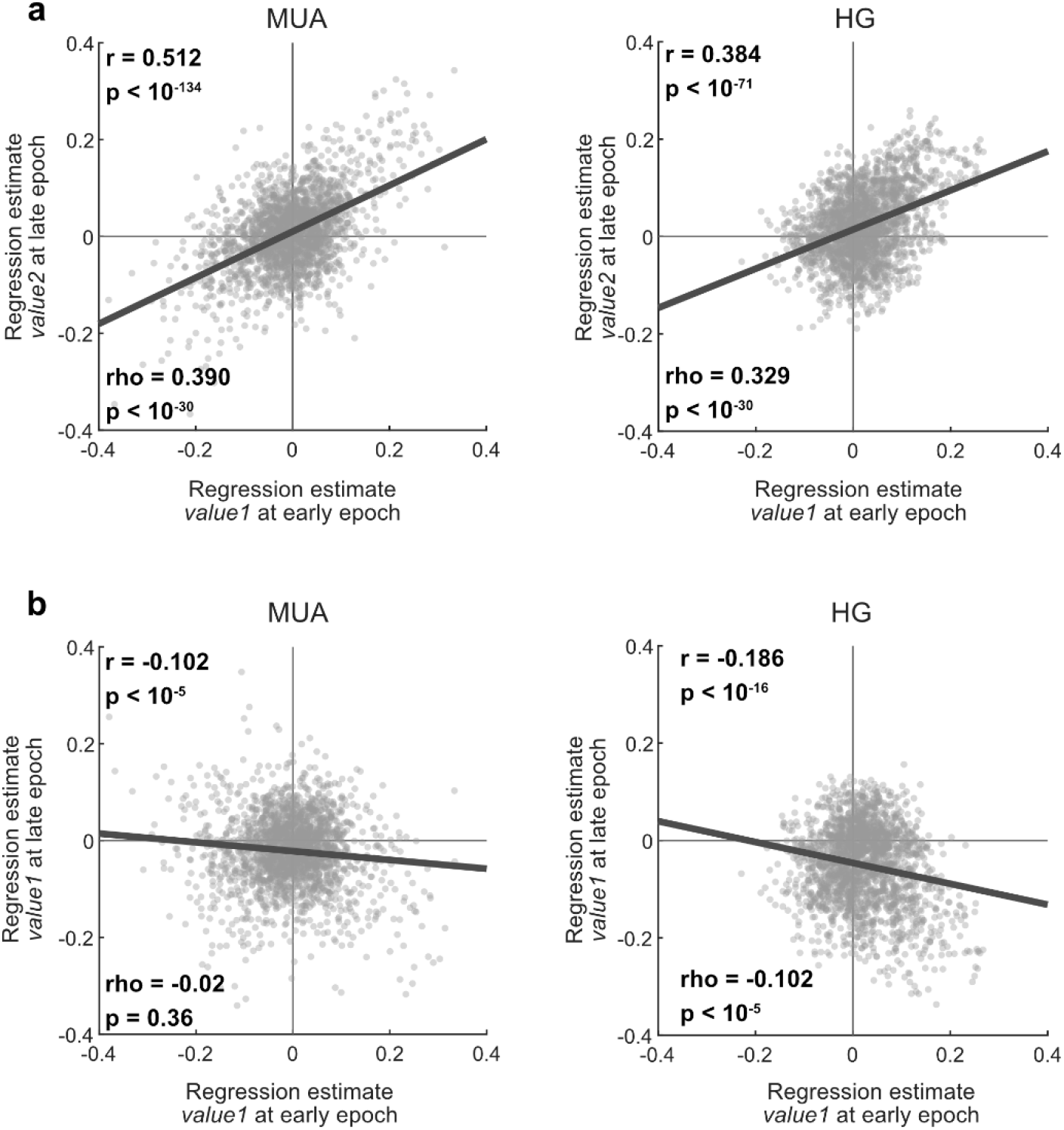
(**a**)Relationship between regression estimates of *value1* atier the first target and *value2* atier the second target. (**b**) Relationship between *value 1* atier the first target and *value1* atier the second target. Early and late epoch indicate 200-400ms atier viewing the first and second targets, respectively (shaded regions in Fig. 4). Pearson’s and Spearman’s correlation coefficients are indicated by ‘r’ and ‘rho’, respectively.

**Supp. Fig. 6.**
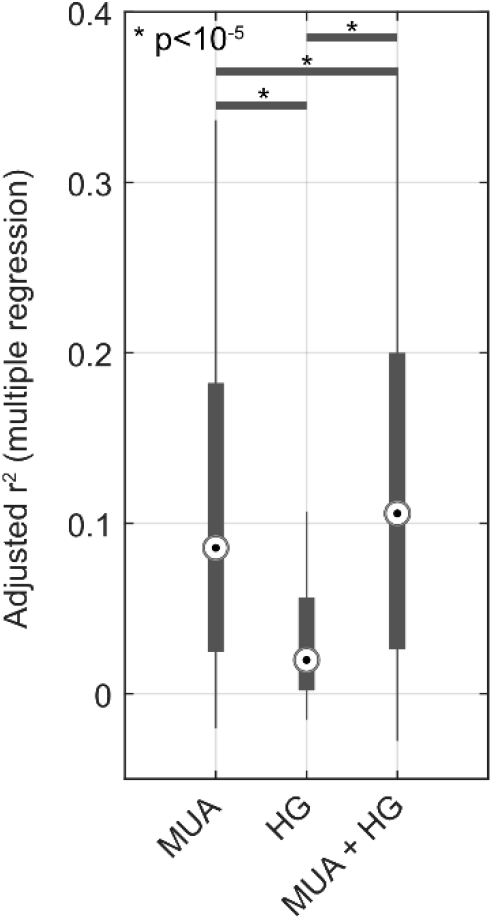
Variance explained in value using models that pool signals across channels in a probe (supporting Fig. 8). Each box plot indicates explained variance in *value1* using pooled MUA only (Eq. 3), pooled HG only (Eq. 3), or both signals pooled together (Eq. 5) for all probes (n=86). P-values give the results of a paired t-test across probes. Data are from 200-400ms atier viewing the first target (early epoch in Fig. 4a).

## Notes

### Competing Interest Statement

The authors have declared no competing interest.

